# Inka2, a novel Pak4 inhibitor, regulates actin dynamics in neuronal development

**DOI:** 10.1101/2022.03.17.484704

**Authors:** Seiya Yamada, Tomoya Mizukoshi, Akinori Tokunaga, Shin-ichi Sakakibara

## Abstract

The actin filament is a fundamental part of the cytoskeleton defining cell morphology and regulating various physiological processes, including filopodia formation and dendritic spinogenesis of neurons. Serine/threonine-protein kinase Pak4, an essential effector, links Rho GTPases to control actin polymerization. Previously, we identified the *Inka2* gene, a novel mammalian protein exhibiting sequence similarity to *Inka1*, which serves as a possible inhibitor for Pak4. Although *Inka2* is dominantly expressed in the nervous system and involved in focal-adhesion dynamics, its molecular role remains unclear. Here, we found that Inka2-iBox directly binds to Pak4 catalytic domain to suppress actin polymerization. Inka2 promoted actin depolymerization and inhibited the formation of cellular protrusion caused by Pak4 activation. We further generated the conditional knockout mice of the *Inka2* gene. The beta-galactosidase reporter indicated the preferential *Inka2* expression in the dorsal forebrain neurons. Cortical pyramidal neurons of *Inka2*^-/-^ mice exhibited decreased density and aberrant morphology of dendritic spines with marked activation/phosphorylation of downstream molecules of Pak4 signal cascade, including LIMK and cofilin. These results uncovered the unexpected function of endogenous *Pak4* inhibitor in neurons. Unlike *Inka1, Inka2* is a critical mediator for actin reorganization required for dendritic spine development.

## Introduction

The actin filament (F-actin) is a fundamental part of the cytoskeleton regulating various physiological cellular processes involving the membrane dynamics, such as motility and cell morphology. For instance, the barbed ends of polymerizing F-actin push the plasma membrane to form cellular protrusions. The Rho family GTPases, including Rac and Cdc42, regulate this highly orchestrated bidirectional process of actin polymerization and depolymerization through several signaling cascades (del Mar Maldonado and Dharmawardhane, 2018). p-21 activated kinase (Pak)/LIM kinases (LIMK)/Cofilin signaling cascade is the central axis regulating F-actin dynamics (Dan et al., 2001; Meng et al., 2002). The Pak family, an evolutionarily conserved serine/threonine-protein kinase, phosphorylates and activates LIMK upon Rac and Cdc42 signaling. Then, the activated LIMK phosphorylates and inactivates the actin-binding protein cofilin, which severs and depolymerizes the F-actin, resulting in the formation and stabilization of the actin cytoskeleton. Pak family consists of six isoforms subdivided into two groups based on the domain architecture: group I (Pak 1, 2, and 3) and group II (Pak 4, 5, and 6), both having distinct and overlapping functions. Pak4 is expressed in diverse tissues, and *Pak4* knockout mice are embryonically lethal, suggesting the essential roles of Pak4 in normal development (Qu et al., 2003; Zhang et al., 2022). Previous studies have shown that constitutively active Pak4 mutant, Pak4 (S445N), phosphorylates LIMK and cofilin to induce cell morphology changes by altered actin polymerization of C2C12 and NIH3T3 cells (Baskaran et al., 2012; Dan et al., 2001; Qu et al., 2001). In contrast, *Pak4* knockdown induces the cell rounding attributed to the actin depolymerization in tumor cell lines H1229 and HCT116 cells (Tabusa et al., 2013; Zhao et al., 2017), suggesting that the control of the fine morphology of cells requires strict regulation of Pak4 activity.

In our previous study, we identified *Inka2* (*Fam212b*) as an evolutionarily conserved gene in various vertebrates and preferentially expressed in the CNS (Iwasaki et al., 2015). Inferring from the partial similarity of the predicted protein structure, *Inka2* and *Inka1* genes seem to constitute the Inka family, characterized by a short conserved motif called “Inka-box (iBox).” Nevertheless, the similarity in function between these two genes remains uncertain because they do not show any sequence similarity besides iBox (Iwasaki et al., 2015). Genetic ablation of *Inka1* in mice results in embryonic exencephaly due to failure of neural tube closure (Reid et al., 2010). Inka1 directly binds the catalytic domain of Pak4 *in vitro* via iBox, repressing the kinase activity of Pak4 (Baskaran et al., 2015). *Inka2* gene is transcriptionally controlled by a tumor suppressor and transcription factor *p53* and acts as a potent inhibitor for cancer cell growth. In addition, the direct interaction of Pak4 with Inka2-iBox is attributable to the anti-tumor effect of *Inka2* (Liu et al., 2019); however, the physiological function of Inka2 in neurons remains unclear.

In the mammalian central nervous system (CNS) development, immature neurons generated from neural stem/progenitor cells (NSPCs) migrate into the cortical plate through sequential morphological changes and polarity formation (Hansen et al., 2017). In the postnatal period, pyramidal neurons arriving at the cortical plate begin to form dendritic spines (Basu and Lamprecht, 2018; Hansen et al., 2017; Nishiyama, 2019). Dendritic spines are actin-based dynamic small protrusions from dendritic shafts that continuously form, change shape, and eliminate throughout life (Nishiyama, 2019). Dendritic spines serve as a storage site for synaptic strength and help transmit electrical signals to the neuron cell body. The appropriate density and morphological type of dendritic spines are crucial for many physiological processes. For example, changes in spine morphology affect the efficacy of memory formation and learning (Frank et al., 2018).

In addition, the loss or malformation of spines is frequently associated with various psychiatric disorders, including schizophrenia, and autism spectrum disorder, including fragile X syndrome (FXS) caused by the mutation in the *fragile X mental retardation 1* (*FMR1*) gene (Davis and Broadie, 2017). Despite the critical role of actin dynamics in spinogenesis, the mechanisms regulating F-actin polymerization and depolymerization in the dendritic spine remain lesser-known. Several studies have revealed that Pak/LIMK/Cofilin signaling is related to dendritic spine morphology in the pyramidal neurons (Meng et al., 2002; Zhang et al., 2005). Increased Pak signaling induces the formation of dendritic spines and the unfunctional dendritic protrusions (Hayashi et al., 2007; Zhang et al., 2005). *FMR1* knockout (KO) mouse, a mouse model of FXS, displays the formation of immature and excessive dendritic spines on cortical neurons similar to the symptoms of FXS in humans (Davis and Broadie, 2017). The administration of a Pak inhibitor rescues the dendritic spine defect in *FMR1* KO mice and ameliorates the autism-like behavioral phenotypes (Dolan et al., 2013). This finding indicates the relevance of the intrinsic system that negatively regulates Pak signaling to prevent abnormal spine formation, facilitating healthy brain development. However, such endogenous Pak inhibitor has not yet been identified in neurons.

Here, we report for the first time that Inka2 functions as the endogenous inhibitor of Pak4 in neurons. Inka2-mediated repression of Pak4/LIMK/cofilin signaling cascade induces the actin depolymerization and impacts the dendritic spine formation.

## Results

### Inka2 causes cell rounding in iBox-dependent manner

To directly assess the role of Inka2 *in vivo*, we first overexpressed *Inka2* in the developing cerebral cortex using *in utero* electroporation technique. Full-length *Inka2* fused with an enhanced green fluorescent protein (EGFP) or control EGFP were electroporated into NSPCs of the E14.5 embryonic neocortex. After 24 h of electroporation, most control EGFP^+^ migrating cells toward the cortical plate had neuronal processes and exhibited multipolar or bipolar morphology (Figure 1A). In contrast, EGFP-Inka2^+^ cells showed a striking change in morphology with neurite retraction and cell rounding (Figure 1A). To assess the association of EGFP-Inka2 with F-actin, the electroporated brain sections were co-stained with phalloidin, a highly selective bicyclic peptide used for staining F-actin. As shown in Figure 1B, overexpressed EGFP-Inka2 was frequently localized on the phalloidin^+^ cell cortex (also known as the actin cortex), which is a specialized layer comprising the F-actin-rich network on the inner face of the cell membrane (Baskaran et al., 2021; Stricker et al., 2010). Physiologically, the onset of apoptosis or cell division is characterized by morphological changes, including cell rounding. However, DNA staining with Hoechst dye showed little or no condensation of chromosomes in *Inka2*-transfected cells, indicating that these cells were not in the mitotic phase (Figure 1B). Moreover, terminal deoxynucleotidyl transferase dUTP nick end labeling (TUNEL) staining of the E15.5 brain sections revealed that most EGFP-Inka2^+^ cells were not engaged in apoptotic states (Figure 1C, D). These results implied an association between Inka2 and F-actin *in vivo*, suggesting that the cell rounding phenotype induced by Inka2 expression resulted from an aberrant reorganization of F-actin.

**Figure 1.**
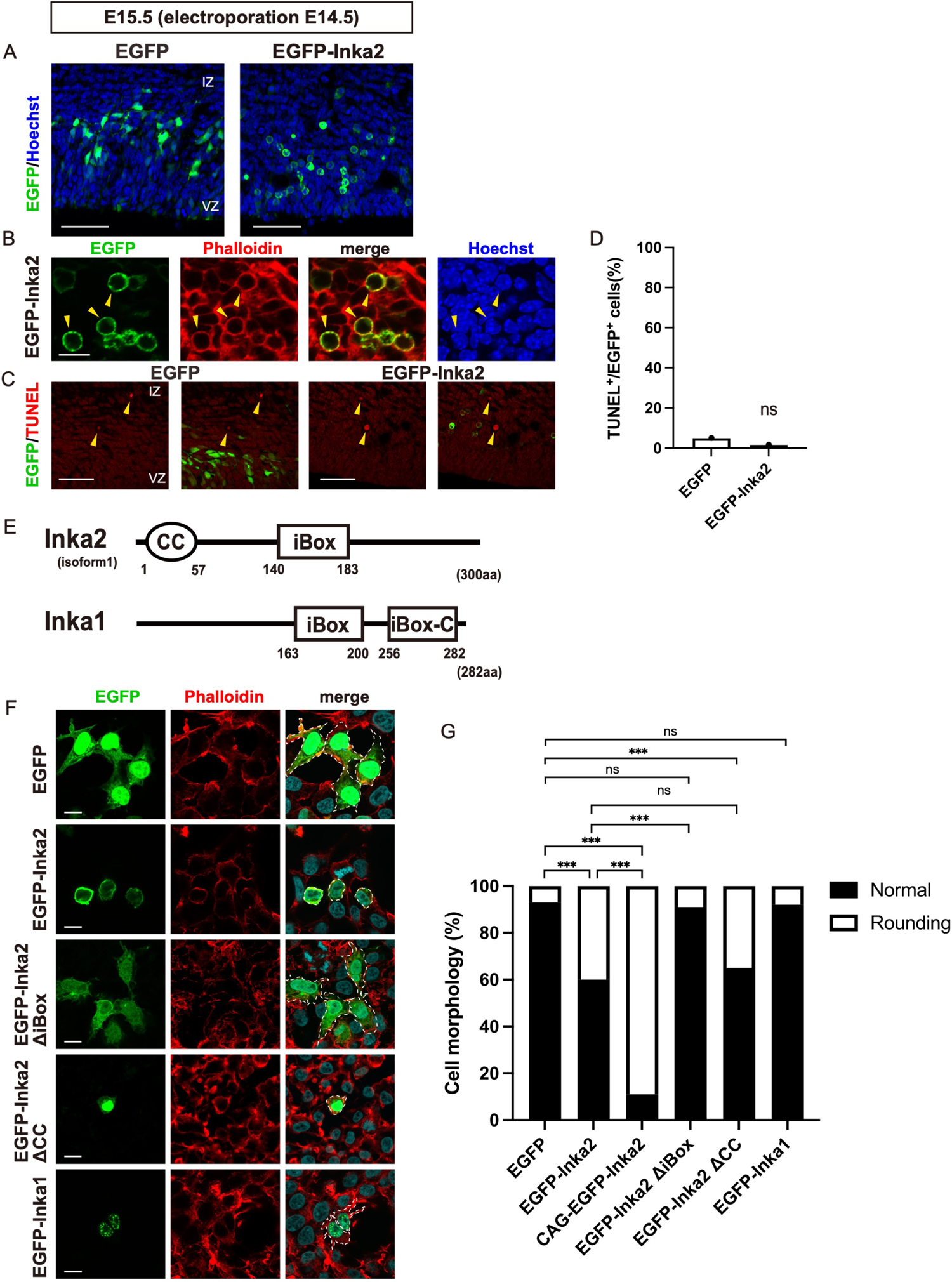
Inka2 overexpression leads the cell rounding *in vivo* and *in vitro*. (A–D) *In utero* electroporation of control CAG-EGFP or CAG-EGFP-Inka2 was performed on E14.5, and the neocortex was analyzed at E15.5. Nuclei were counterstained with Hoechst dye (blue). (B) Brain sections electroporated with CAG-EGFP-Inka2 were stained with phalloidin (red). Arrowheads denote the rounding cells. (C) Apoptotic cells were detected by TUNEL staining (red) at E15.5. Arrowheads indicate TUNEL^+^ cells. (D) Quantified comparison of the EGFP^+^ TUNEL^+^ apoptotic cells. ns, not significant; chi-square test. n = 60 for EGFP^+^ cells; n = 59 for EGFP-Inka2^+^ cells. (E) Domain structure of mouse Inka2 and Inka1. Oval and rectangle indicate coiled-coil (CC) and Inka-box (iBox) domain, respectively. Inka1 has two iBox domains. VZ, ventricular zone; IZ, intermediate zone. (F, G) HEK293T cells were transfected with control EGFP, EGFP-Inka2, CAG-EGFP-Inka2, EGFP-Inka2ΔiBox, EGFP-Inka2ΔCC, or EGFP-Inka1.F-actin was stained with phalloidin (red). The areas surrounded by dashed lines exhibit the cellular morphology. (G) Quantified comparison of the effect of Inka2 on cell morphology. The stacked bar chart shows the percentages of cells that exhibit a normal or rounding morphology. ns, not significant; ***, P < 0.001; chi-square tests with Holm–Bonferroni correction. EGFP, n = 138 cells; EGFP-Inka2, n = 127 cells; CAG-EGFP-Inka2, n = 95 cells; EGFP-Inka2ΔiBox, n = 119 cells; EGFP-Inka2ΔCC, n = 155 cells; EGFP-Inka1, n = 214 cells. Unless otherwise specified in (F) and (G), EGFP, EGFP-Inka1, EGFP-Inka2, and EGFP-Inka2 deletion mutants represent the expression constructs driven by the CMV promoter. Scale bars, 50 μm in (A) and (C), and 10 μm in (B) and (F).

A homology search against the protein databases, including BLASTP, revealed that Inka2 shared a sequence similarity (approximately 70%) with Inka1 in the iBox region (amino acid residues 140–183) (Figure 1E). The iBox contains the tripeptide PLV sequence in common with the substrate-docking site of Pak4 (Baskaran et al., 2015). Although the physiological significance of the iBox has not been fully understood, the Inka1 protein was shown to interact with Pak4 via the iBox domain (Baskaran et al., 2015) physically. Of note, an *in silico* analysis using COILS (Lupas et al., 1991) and Paircoil2 (McDonnell et al., 2006) algorithms predicted the presence of coiled-coil (*CC*) domain (residues 1–57) in the N-terminal region of Inka2 (Figure 1E). *CC* is a helical structure of α-helices found in various proteins, including transcription factors, cytoskeletons, and motor proteins, and is known to be involved in protein– protein interaction or function of several classes of fibrous structural proteins (Rose and Meier, 2004). Considering the low sequence similarity outside the iBox and the presence of *CC, Inka2 and Inka1* genes likely have discrete molecular properties.

To evaluate the effect of Inka2 and Inka1 on cell morphology, EGFP-Inka2 or EGFP-Inka1 was introduced *in vitro* in HEK293T cells. In control EGFP^+^ cells, phalloidin^+^ F-actin was localized at the cell periphery, and most cells showed standard amorphous shape (normal shape, 93.5%; rounding shape, 6.5%) (Figure 1F, G). EGFP-Inka2 expression significantly elevated a cell population with abnormal morphology frequently characterized by cell rounding (normal, 59.8%; rounding, 40.2%). EGFP-Inka2, dominantly localized to the peripheral cell cortex, was positive for phalloidin, as *in vivo* overexpression in the embryonic brain (Figure 1B). Such a cell rounding phenotype was hardly caused by the EGFP-Inka1 overexpression (normal, 92.1%; rounding, 7.9%) (Figure 1F, G). Moreover, EGFP-Inka1 was localized to the granule structures in the nucleus (Figure 1F), which is consistent with the previous study (Baskaran et al., 2015). These observations suggested a unique function of Inka2 in actin reorganization controlling the cell morphology. To identify the Inka2 domain having a central role in the morphological change of cells, the deletion constructs lacking the CC domain (EGFP-Inka2ΔCC), or iBox domain (EGFP-Inka2ΔiBox) were transfected into HEK293T cells. While EGFP-Inka2ΔCC caused cell rounding (normal, 65.2%; rounding, 34.8%), EGFP-Inka2ΔiBox overexpression abolished this morphological phenotype (normal, 90.8%; rounding, 9.2%) (Figure 1F, G), indicating the importance of Inka2 iBox domain in the cell rounding phenotype.

### Inka2 binds Pak4 via iBox and regulates Pak4 signaling to control cell morphology

Both human Inka2 and Inka1 interact *in vitro* with Pak4 via iBox (Baskaran et al., 2015; Liu et al., 2019), although whether Inka family members were implicated in the dynamics of F-actin through the modulation of Pak4 activity remains unknown. First, the direct binding of Inka1 or Inka2 with Pak4 was confirmed by co-immunoprecipitation (Co-IP) using HEK293T cells expressing Flag-Pak4 and EGFP-Inka1 or EGFP-Inka2 (Figure 2A). Inka2ΔiBox showed no binding with Pak4, indicating the iBox-dependent interaction between Inka2 and Pak4 (Figure 2A). Next, we investigated whether the cell rounding phenotype induced by Inka2 depends on Pak4 activity. Figure 2B shows the individual cell overexpressing EGFP-Inka1 or Inka2, confirming that Inka2, but not Inka1, induced cell rounding. Substantially, this Inka2 phenotype was rescued by the co-expression with Pak4. Many EGFP-Inka2 cells co-expressing with Flag-Pak4 exhibited normal morphology (Figure 2C). Consistent with the previous result (Qu et al., 2001), the overexpression of Pak4 alone never induced a significant morphological change in cells (Figure 2D–F). Additionally, we observed that the occurrence ratio of cells rounding up was proportional to the expression level of EGFP-Inka2. As shown in Figure 2D, the frequency of rounding cells was greatly enhanced by Inka2 expression driven with CAG promoter (normal, 10.5%; rounding, 89.5%), which is a more potent synthetic promoter than the cytomegalovirus (CMV) early enhancer-promoter. Again, Flag-Pak4 co-expression significantly rescued the cell rounding phenotype caused by CAG-EGFP-Inka2 (normal, 66.7%; rounding, 33.3%) (Figure 2C, D). We frequently observed the colocalization of EGFP-Inka2 with Flag-Pak4 in the cell cortex, where phalloidin^+^ F-actin was enriched (Figure 2E, F). In contrast, Pak4 co-expression did not affect the morphology of EGFP-Inka1 cells (normal, 88.3%; rounding, 11.7%) (Figure 2C, D) nor cells overexpressing EGFP-Inka2ΔiBox (Figure 2E–F). These results indicated that the cell rounding is a specific phenotype to Inka2 and that the interaction between Inka2-iBox and Pak4 regulates the cell shape accompanied by actin reorganization. Meanwhile, we often observed the change in subcellular localization of EGFP-Inka1 into cytosol moving from the nucleus upon Pak4 co-expression (Figure 2F bottom panels, compared to Figure 1F). Consistent with our observation, a previous study showed that co-expression with Pak4 causes the translocation of Inka1 protein into the cytosol (Baskaran et al., 2015).

**Figure 2.**
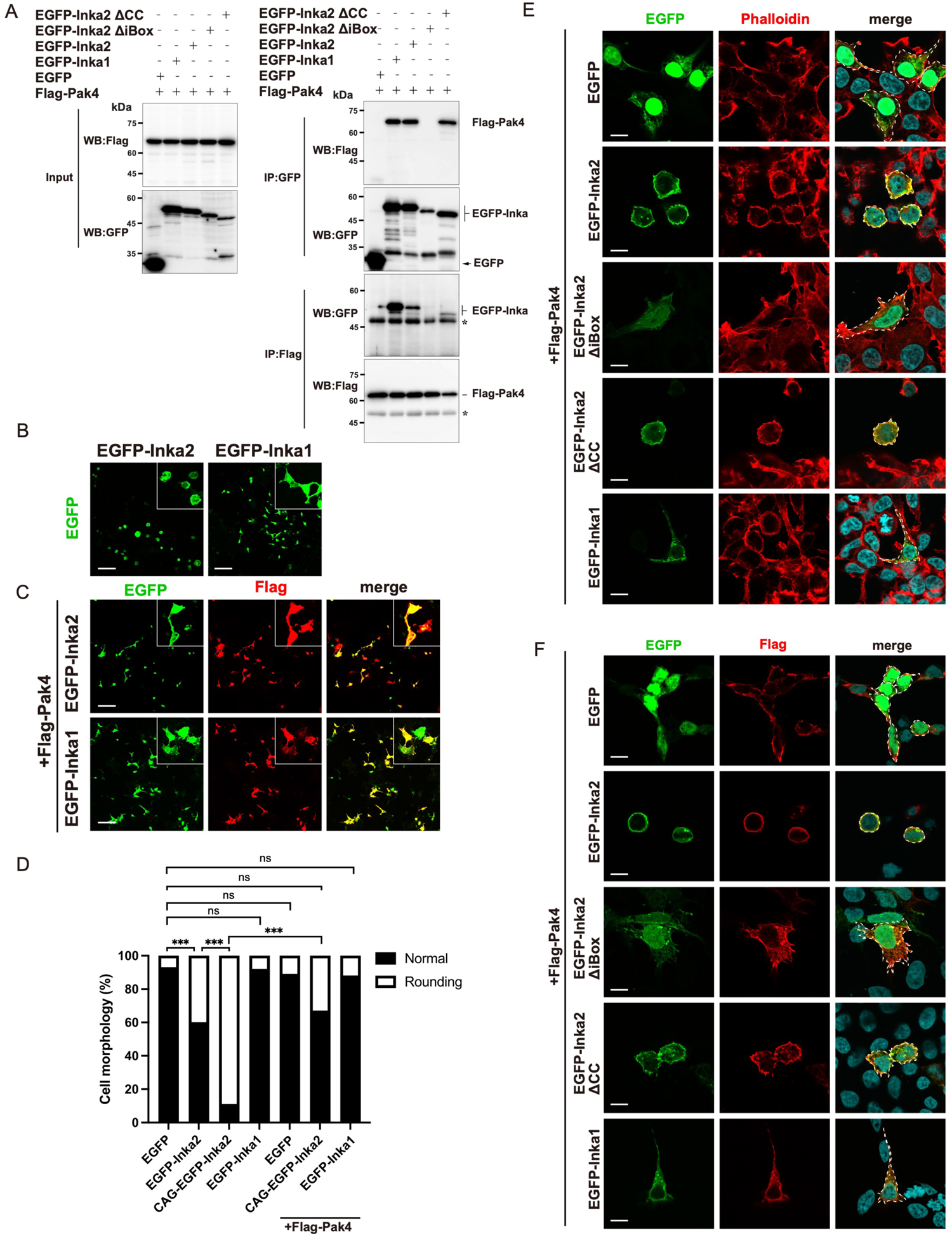
Inka2-iBox interacts with Pak4 to regulate actin reorganization. (A) Co-IP showing the interaction between Inka2-iBox and Pak4. HEK293 cells expressing Flag-Pak4 with EGFP, EGFP-Inka2, EGFP-Inka2ΔiBox, EGFP-Inka2ΔCC, or EGFP-Inka1 were subjected to immunoprecipitation with an anti-GFP or anti-Flag antibodies, followed by immunoblotting with anti-GFP or anti-Flag antibodies (right panels). Left panels showed the input samples, confirming the expression of each construct. (B–F) HEK293T cells were transfected with Flag-Pak4 with EGFP, EGFP-Inka2, EGFP-Inka2, or EGFP-Inka1 (B, C). Insets showed a magnified view of the individual cell morphology. (C, F) Cells were immunostained with an anti-Flag antibody (red). (E) F-actin was stained with phalloidin (red). (E, F) The areas surrounded by dashed lines exhibit the cellular morphology. (D) Quantified comparison of the effect of Inka2 on cell morphology. The stacked bar chart shows the percentages of cells that exhibit a normal or rounding. To estimate the effect of Flag-Pak4 on cell morphology, data pertaining to EGFP, EGFP-Inka2, and -Inka1 in Figure 1G were shown again, which were measured in parallel in the absence of Flag-Pak4. ns, not significant; ***, P < 0.001; chi-square tests with Holm–Bonferroni correction. EGFP, n = 138 cells; EGFP-Inka2, n = 127 cells; CAG-EGFP-Inka2, n = 95 cells; EGFP-Inka1, n = 214 cells; EGFP+Pak4, n = 91 cells; CAG-EGFP-Inka2+Pak4, n = 114 cells; EGFP-Inka1+Pak4, n = 111 cells. Unless otherwise specified, Flag-Pak4, EGFP, EGFP-Inka1, EGFP-Inka2, and EGFP-Inka2 deletion mutants represent the expression constructs driven by the CMV promoter. Scale bars, 100 μm in (B) and (C), and 10 μm in (E) and (F).

Under the physiological condition, intracellular Pak4 activity is autoinhibited and hardly affects the cell morphology in a steady-state (Baskaran et al., 2012; Dan et al., 2001; Qu et al., 2001). To clarify the effect of Inka2 on Pak4 signaling, we evaluated the morphological changes in HeLa cells expressing the constitutive active mutant of Pak4 (Pak4cat) (286–591 aa), in which the autoinhibitory domain was deleted. Without the Pak4cat expression, HeLa cells exhibited an epithelial cell-like morphology with irregular polygonal shapes without long fine processes. HeLa cells transfected with EGFP or Flag empty vector showed normal morphology cells with smooth contour (normal polygonal cells, 98.0%; cells with long processes, 2.0%) (Figure 3). We discovered that Flag-Pak4cat expression drastically altered the shape of HeLa cells to extend several fine long processes, as previously described (Dan et al., 2002), which were positive for phalloidin in a dose-dependent manner of Pak4cat (population of cells with long processes, 13.7% [0.1μg of Pak4cat]; 25.4% [0.5μg]; 46.9% [1.5μg]) (Figure 3B). Concomitant expression of EGFP-Inka2 diminished these abnormal cell morphology induced by Pak4cat (Figure 3A, B). Unlike the case of HEK293T cells, the overexpression of EGFP-Inka2 alone caused little or no morphological change in HeLa cells (Figures 3 and S1). Failure of the phenotype rescue by EGFP-Inka2ΔiBox further supported the idea that Inka2 represses Pak4 activity via iBox to control actin reorganization (Figure 3C). In contrast, consistently with the previous study (Baskaran et al., 2015), co-expression of Pak4cat and EGFP-Inka1 resulted in the formation of intracellular protein crystal composed of Pak4cat and EGFP-Inka1 (Figure S2); thus, we could not assess the effect of Inka1 on Pak4 activity in our assay.

**Figure 3.**
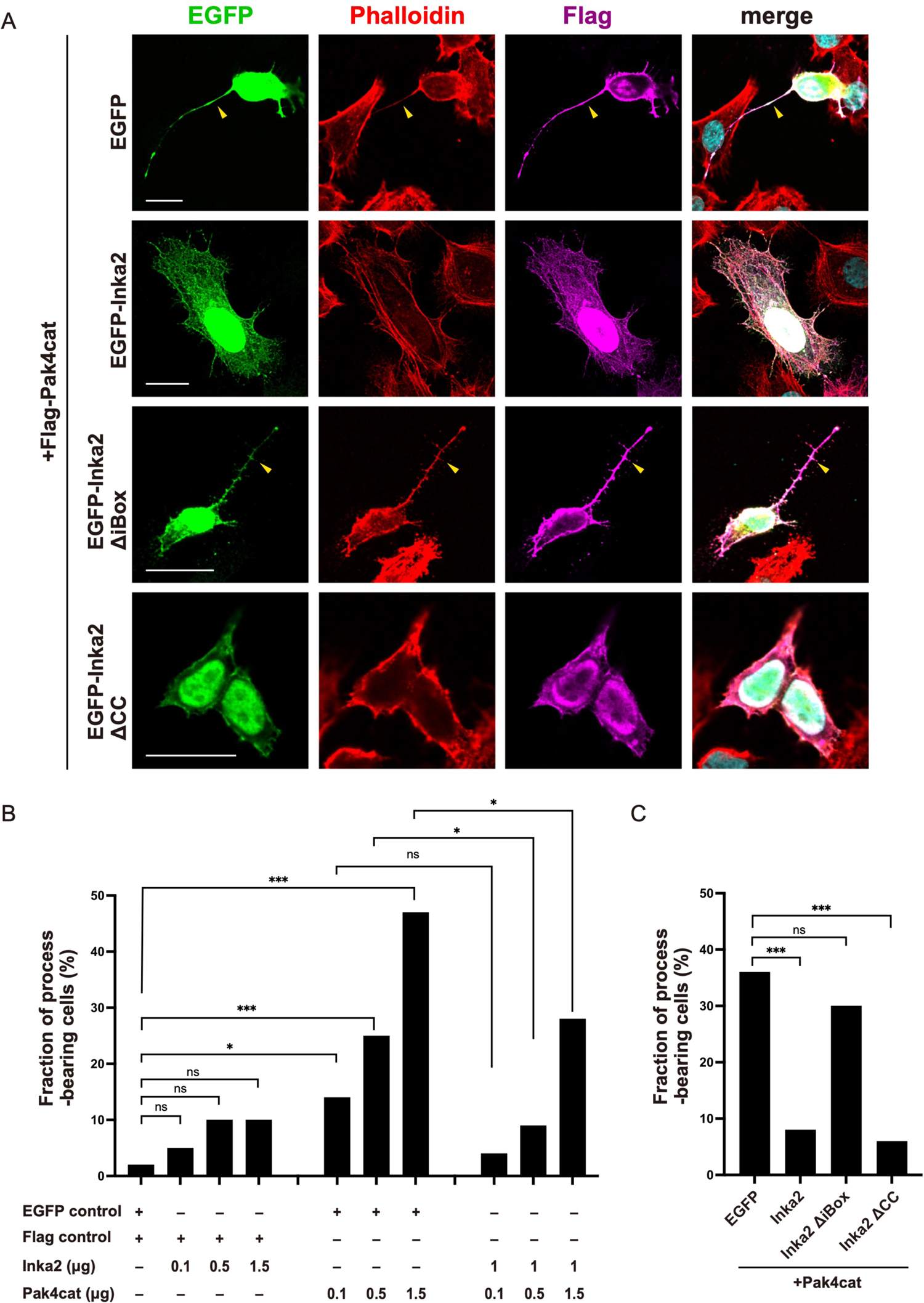
Inka2 inhibits Pak4-dependent morphological change of cells. (A–C) HeLa cells were transfected with EGFP, EGFP-Inka2 (0.1 μg–1.5 μg), EGFP-Inka2ΔiBox, or EGFP-Inka2ΔCC along with Flag-control or Flag-Pak4cat (0.1 μg–1.5 μg). Cells were immunostained with an anti-Flag antibody (magenta). F-actin was visualized by phalloidin (red). Arrowheads indicate the abnormally extended cell processes. (B) Quantified comparison of the effect of Inka2 (0.1 μg–1.5 μg) and Pak4cat (0.1–1.5 μg) on cell morphology. (C) Morphological effect of Inka2, Inka2ΔiBox, or EGFP-Inka2ΔCC (1 μg) on the HeLa cells expressing Pak4cat (1 μg). The stacked bar chart shows the percentages of cells bearing the fine long processes. All expression constructs were driven by the CMV promoter. ns, not significant; *, P < 0.05; ***, P < 0.001; chi-square tests with Holm–Bonferroni correction. n = 65–150 cells in each condition. Scale bars, 20 μm in (A).

### Inka2 elicits depolymerization of actin filaments as Pak4 specific inhibitor

The inhibitory effect of Inka2 for actin polymerization was investigated using an *in vitro* cell-free reconstitution assay. Bacterially-expressed and -purified recombinant proteins of GST-Inka2-iBox or His-Pak4cat (Figure 4A) were added to the HEK293 cell lysate. Then the filamentous F-actin pool and the free monomeric globular actin (G-actin) pool were fractionated and quantified using analytical differential ultracentrifugation followed by immunoblotting using anti-β actin antibody (Figure 4B). Compared with control GST, His-Pak4cat alone did not cause a significant fluctuation of the F/G-actin ratio (F-actin content in GST control as 1.0; GST + Pak4cat, 1.07 ± 0.04). In contrast, simultaneous incubation of Inka2-iBox with Pak4cat substantially decreased the F/G-actin ratio (GST-Inka2-iBox + Pak4cat; 0.68 ± 0.05) (Figure 4C), indicating that Inka2 induced destabilization and depolymerization of F-actin. We further determined the F-actin and G-actin contents in HEK293T cells overexpressing EGFP-Inka2 or EGFP-Inka1 and observed that Inka2 slightly decreased the F-actin/G-actin pool ratio (Figure S3A, B). Pak4 overexpression partially restored the depolymerization caused by Inka2 (Figure S3B). Compared to Inka2, EGFP-Inka1 exhibited no or a slight effect on actin depolymerization (Figure S3B).

**Figure 4.**
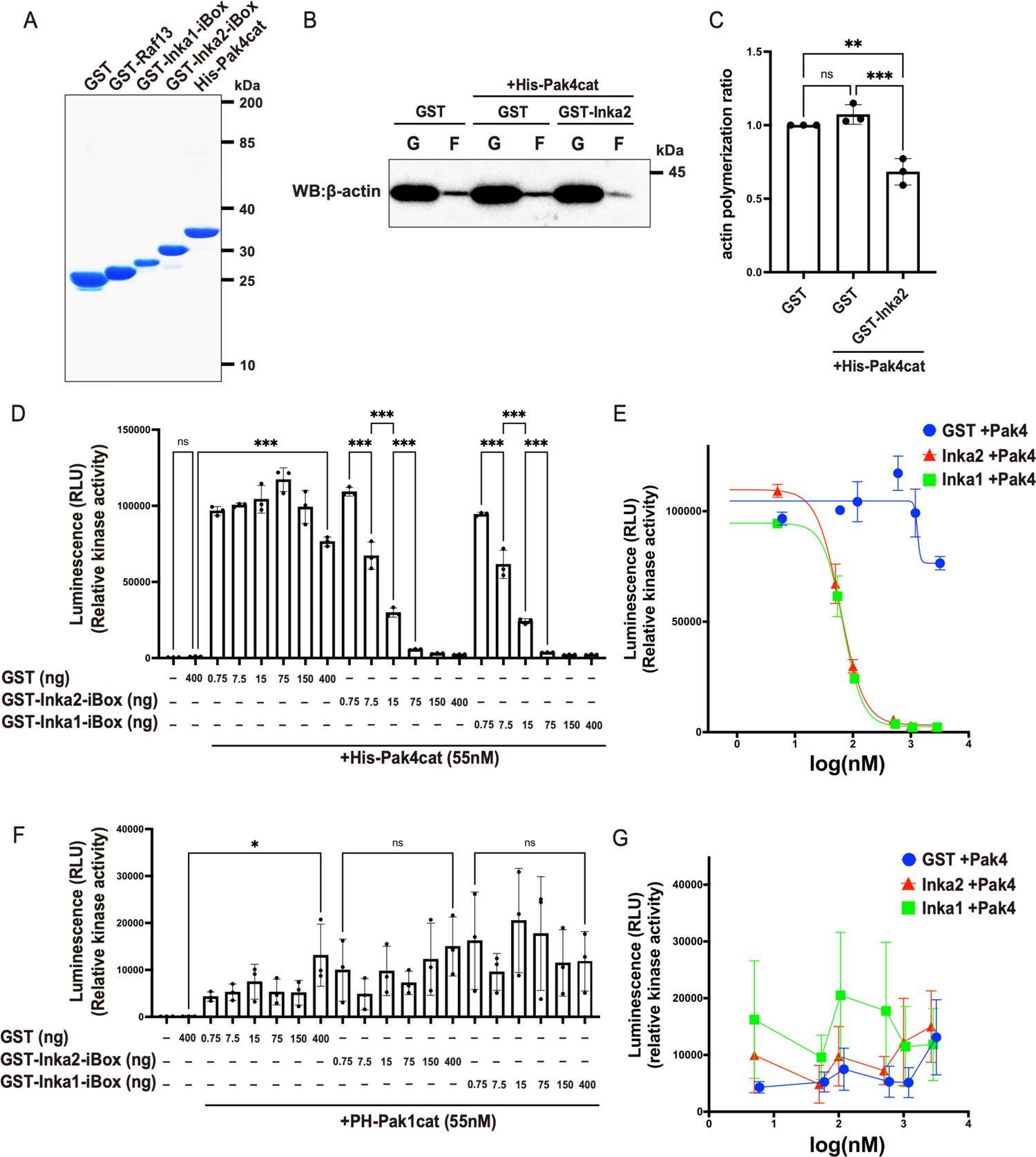
Inka2 inhibits Pak4 activity and depolymerizes F-actin. (A) Purified recombinant proteins of GST, GST-Raf13, GST-Inka1-iBox, GST-Inka2-iBox, and His-Pak4cat. SDS-PAGE and Coomassie brilliant blue staining confirmed the quality of each protein. (B, C) Cell lysate prepared from HEK293T cells was incubated with the purified proteins of GST or GST-Inka2-iBox together with His-Pak4cat. After incubation, F-actin and G-actin fractions were separated and quantified by immunoblotting with an anti-β-actin antibody. (C) Quantified analysis of actin polymerization (F-actin/G-actin) ratio of three independent experiments. ns, not significant, *, P < 0.05; One-way ANOVA. Holm–Sidak’s multiple comparisons test. (D) *In vitro* kinase assay using His-PAK4cat with GST, GST-Inka1-iBox, and GST-Inka2-iBox. Pak4 activity was assessed using AKT1 peptide substrate and represented as the luciferase-based luminescence (RLU). Three independent experiments were performed. ns, not significant; ***, P < 0.001; One-way ANOVA. Holm–Sidak’s multiple comparisons test. (E) Inhibition profile of GST, GST-Inka1-iBox, and GST-Inka2-iBox on Pak4 activity. The IC_50_ values (Inka1, 67.7 nM; Inka2, 60.3 nM) were calculated from the Sigmoid curve. (F) *In vitro* kinase assay using Pros2-His-PAK1cat (PH-Pak1cat) with GST, GST-Inka1-iBox, and GST-Inka2-iBox. Pak1 activity was determined as above. Three independent experiments were performed. ns, not significant; *, P < 0.05; One-way ANOVA. Holm–Sidak’s multiple comparisons test. (G) Inhibition profile of GST, GST-Inka1-iBox, and GST-Inka2-iBox on Pak1 activity.

To examine whether the direct binding of Inka2 and Pak4 causes inhibition of Pak4 activity, we next performed *in vitro* kinase assay of Pak4 with or without Inka1-iBox or Inka2-iBox (Figure 4A). The Ser/Thr kinase activity of purified His-Pak4cat protein was measured in the presence of recombinant purified protein GST, GST-Inka1-iBox, or GST-Inka2-iBox using synthetic peptide of AKT1 or GST-Raf13 as the substrate. Inka2-iBox directly inhibited the Pak4 activity in a dose-dependent manner (Figure 4D). This inhibitory potency of Inka2 was almost identical to that of Inka1 (Figure 4D), as the half-maximal inhibitory concentration (IC_50_) calculated based on the dose-response curve was estimated as 60.3 nM for GST-Inka2-iBox and 67.7 nM for GST-Inka1-iBox (Figure 4E). Members of the Ser/Thr kinase Pak family recognize distinct substrate specificity and play different functional roles. Therefore, we surveyed the Inka2 inhibitory effect on Pak1 activity, a member of the Pak family in group I. The kinase activity of Pak1 was neither inhibited by Inka2-iBox nor Inka1-iBox (Figure 4F, G). Altogether, these findings indicated that Inka2 specifically inhibits Pak4 activity through direct binding with iBox, directing the depolymerization of F-actin.

### Generation of *Inka2* knockout (KO) mice

Mouse *Inka2* gene is mapped to chromosome 3 and consists of two exons (Figure 5A). To reveal the *in vivo* function of the *Inka2* gene, we set out the generation of *Inka2* KO mice. ES cells carrying *Inka2* KO-first allele (*Inka2^tm1a^*) were obtained from the Knockout Mouse Project (KOMP) and used to produce chimera and *Inka2^tm1a^* heterozygous mutant mice. *Inka2^tm1a^* is a floxed allele with conditional KO potential harboring the β-galactosidase (lacZ) cassette as a gene expression reporter (Figure 5A). We referred *Inka2^tm1a^* allele as *Inka2*^flox^ hereafter.

**Figure 5.**
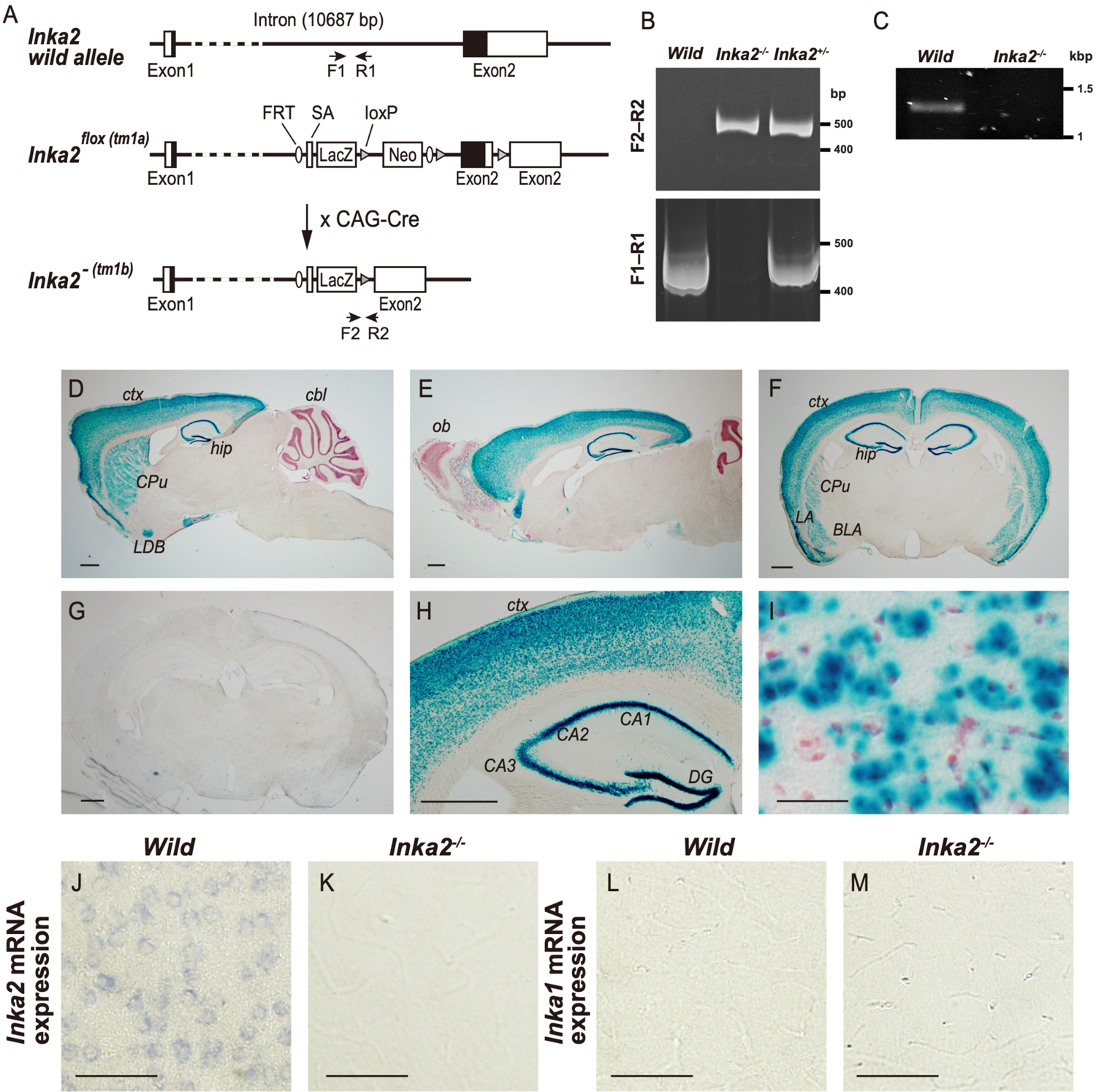
Generation of *Inka2 k*nockout mice. (A) Schematic diagram of the *Inka2* wild-type allele (top), and the conditional allele of *Inka2*^flox^ (*Inka2*^tm1a(KOMP)Wtsi^) (middle). The splicing of exon1 to a lacZ trapping element occurs in the *Inka2^flox^* allele because the lacZ and neo cassette inserted in the *Inka2* intron 1 contains the Engrailed2 splice acceptor (En2SA) and the SV40 polyadenylation sites. *Inka2*^flox^ mice were crossed with the CAG-Cre transgenic mice harboring the CAG-promoter driven Cre recombinase to generate *Inka2*^−^ (*Inka2*^tm1b(KOMP)Wtsi^) null allele (bottom). Upon elimination of the floxed *Inka2* critical exon (exon 2), this allele generates a truncated RNA lacking most of the open reading frame (ORF) of *Inka2*. (B) Filled box, the *Inka2* ORF; Open box, 5′- and 3′-UTR of *Inka2* gene; SA, En2 splicing acceptor sequence; LacZ, β-galactosidase reporter cassette containing IRES and SV40 polyA site; neo, neomycin resistant gene cassette driven by the SV40 promoter; FRT, FLP recombinase target sequence. (C) PCR genotyping from wild, *Inka2*^+/-^, or *Inka2*^-/-^ tail genomic DNA. The specific primers (primer pairs F1‒R1 and F2‒R2 depicted in A) are used to detect wild-type (402 bp amplicon) and *Inka2*^−^ (430 bp amplicon) alleles. PCR products were separated by 5% PAGE. (D) Loss of *Inka2* mRNA expression in *Inka2*^-/-^ brains. The mRNAs isolated from wild-type and *Inka2*^-/-^ adult mouse cerebral cortex were subjected to 3′RACE analysis using a specific primer that flanks the *Inka2* exon 1. Nested PCR was further performed to confirm the specificity of the PCR product. A representative image of agarose gel electrophoresis is shown. (D–I) lacZ reporter activity in adult *Inka2*^flox/+^ brains. X-gal staining of sagittal sections (D, E) or coronal sections (F, G). (F) X-gal staining of wild-type brain. (H) Higher magnification of the cerebral cortex and hippocampus of D. (I) Magnified view of the cerebral cortex layer Ⅴ. Sections were counterstained with neutral red. (J–M) *Inka2* and *Inka1* mRNA expressions in adult mouse cerebral cortex. *In situ* hybridization of wild-type (J, L) and *Inka2*^-/-^ (K, M) cortical layer V with the *Inka2* (J, K) and *Inka1* (L, M) riboprobe. CPu, caudate putamen; ctx, cerebral cortex; hip, hippocampus; cbl, cerebellum; LDB, lateral nucleus of the diagonal band; ob, olfactory bulb; LA, lateral amygdala nucleus; BLA, basolateral amygdala nucleus; cc, corpus callosum; DG, dentate gyrus. Scale bars, 500 μm in (C–G), 20 μm in (H), (I), and 50 μm in (J–M).

First, we analyzed the lacZ expression in *Inka2*^flox/+^ mice. Enzyme activity of β-galactosidase encoded by lacZ is expected to reflect a transcription level of the *Inka2* promoter. Among various tissues, lacZ was predominantly detected in the nervous system. In the adult brain, a high level of β-galactosidase activity was found in neurons of forebrain regions, including the pyramidal neurons in the cerebral cortex, dentate gyrus granular cells and CA1‒CA3 pyramidal neurons in the hippocampus, and caudate-putamen neurons in the basal forebrain, including the nucleus of the diagonal band (Figure 5D–I). In contrast, lacZ activity was undetectable in the caudal brain areas, including the thalamus, midbrain, pons, medulla, and cerebellum (Figure 5C–H), implicating a forebrain-specific transcription of *Inka2* gene promoter. Control wild-type mice showed no lacZ staining (Figure 5F). Consistently, *in situ* hybridization experiment in our previous study showed that *Inka2 mRNA* expression is rapidly upregulated during the postnatal CNS development (Iwasaki et al., 2015). Moreover, large amounts of *Inka2* mRNA were principally detected in neurons in the adult forebrain (Iwasaki et al., 2015). These results imply that the *Inka2* function is associated with the differentiation or maintenance of the forebrain neurons.

To obtain the *Inka2* null (*Inka2^-/-^*) mice, heterozygous *Inka2*^flox/+^ mice were subsequently crossed with the CAG-Cre deleter mice line, which ubiquitously expresses Cre recombinase, thereby allowing the deletion of the loxP-flanked genomic region (*Inka2*^tm1b^ allele) (Figure 5A). *Inka2*^tm1b^ allele lacks the genomic region encompassing exon 2, a critical exon covering over 90% of the coding region of *Inka2* (Figure 5A).

The *Inka2*^tm1b^ allele was denoted as *Inka2*^-^ in the present study. Interbreeding of the heterozygous mutant (*Inka2*^-/+^) mice yielded homozygous mutant (*Inka2*^-/-^) pups with the expected Mendelian ratio, implying that *Inka2* is not essential for viability. Genomic sequencing and genomic polymerase chain reaction (PCR) analysis verified the correct targeted disruption of the *Inka2* gene (Figure 5B). 3′RACE and quantitative reverse transcription (qRT)-PCR analysis confirmed that *Inak2*^-/-^ mice completely abolished the *Inka2* mRNA expression (Figure 5C, 7A, 7B). *In situ* hybridization analysis demonstrated the *Inka2* mRNA expression in neurons residing in the cerebral cortex (Figure 5J) and the hippocampus (Figure S4A) of wild-type mice, but no signal of *Inka2* mRNA was detected in the *Inka2*^-/-^ brain (Figure 5K, S4B). On the other hand, *Inka1* mRNA was undetectable in all brain regions irrespective of the *Inka2* genotype (Figure 5L, M, S4C, D). Q-PCR analysis further confirmed the undetectable level of *Inka1* mRNA in the *Inka2*^-/-^ cerebral cortex and in wild-type primary cortical neurons (Figure 7B). Based on these observations, we concluded that *Inka1* gene is not expressed in the neuronal population and that its expression is not compensatory upregulated in the *Inka2^-/-^* brain.

Nevertheless, we could not demonstrate the absence of *Inka2* protein product in *Inka2*^-/-^ mice brains because there is no anti-Inka2 antibody suitable for the immunoblotting or immunohistochemistry in mouse tissues. We tried to generate Inka2 antibodies using distinct recombinant proteins as immunogens but failed to produce a specific antibody recognizing endogenous Inka2. A difficulty in detecting endogenous Inka2 might be, at least in part, attributable to the translational repression of Inka2, as described below.

### Disruption of *Inka2* induces an aberrant dendritic spine formation

Although homozygous mutant *Inka2*^-/-^ mice showed no gross defects in development, the detailed histological analysis of the brain revealed the unusual morphological features of *Inka2*^-/-^ neurons. *Inka2* KO adult mice frequently exhibited a moderate dilation of the lateral ventricles with thinner corpus callosum (Figure 6A, C, D).

**Figure 6.**
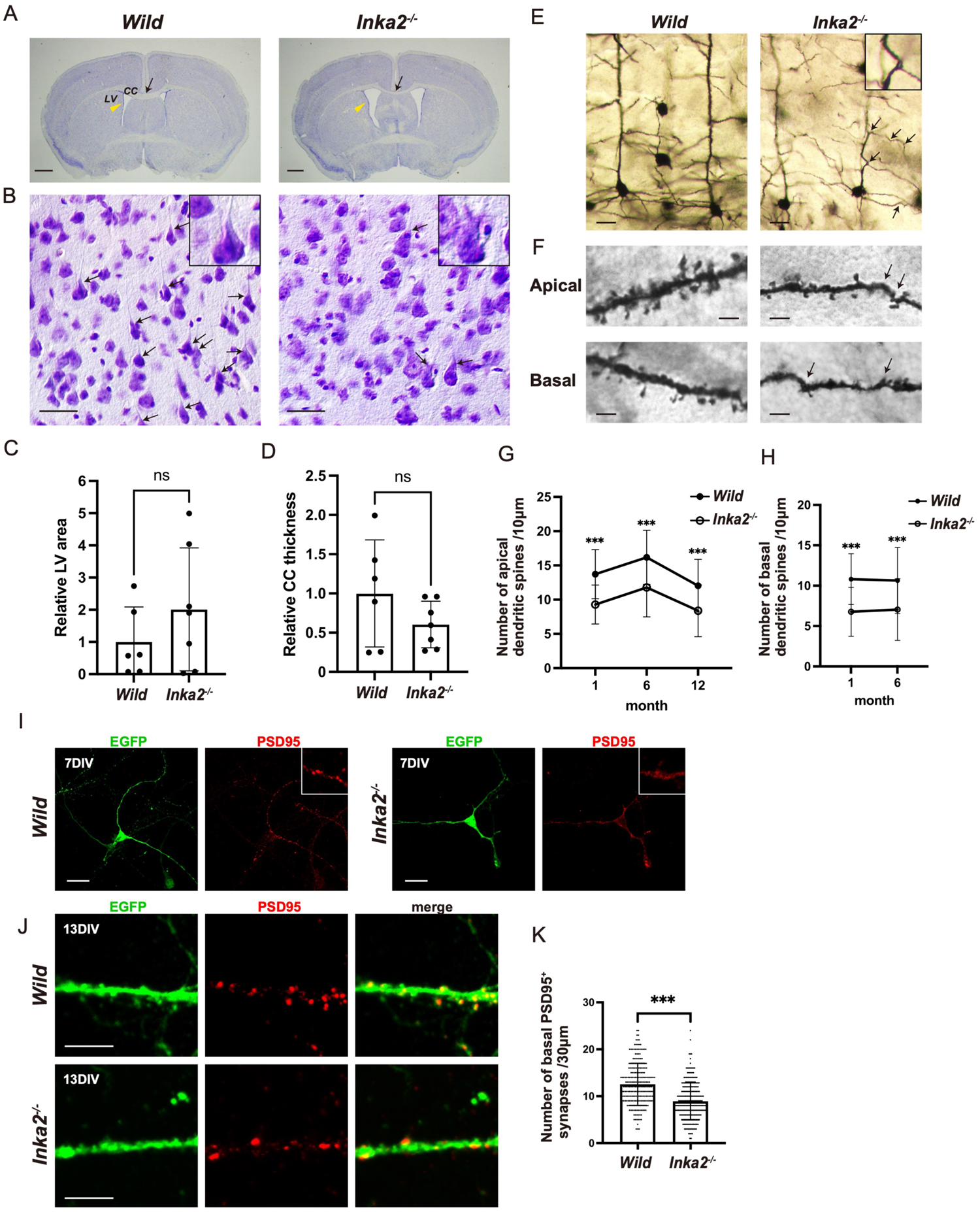
Loss of *Inka2* reduces the dendritic spine formation with deviant morphology. (A–B) Coronal section from wild-type or *Inka2^-/-^* adult brain (Nissl staining). Arrowheads indicate the lateral ventricles (LV). Arrows denote corpus callosum (CC). (B) Higher magnification of layer Ⅴ in the cerebral cortex. Arrows indicate the regular apical dendrites of the cortical neurons. Insets show the pyramidal neurons. (C) Quantification of LV area. n = 7 brains. ns, not significant; Welch’s t-tests. (D) Quantification of the relative thickness of CC. n = 7 brains. ns, not significant; Welch’s t-tests. (E‒H) Golgi silver impregnation staining of cortical layer V in 6-month-old wild-type or *Inka2^-/-^* mouse. Arrowheads indicate abnormal bending of apical or basal dendrites. (F) Higher magnification of dendritic spines of apical and basal dendrites of the pyramidal neuron in layer V. Arrowheads indicate the abnormal bending of dendrite with irregularly distorted morphology. (G, H) The number of spines on the apical dendrite (G) or basal dendrite (H) (per 10 μm length of dendrite) of pyramidal neurons in 1- to 12-month-old wild-type or *Inka2^-/-^* mice. n = 50 different dendrites for each genotype. ***, P < 0.001; One-way ANOVA. Holm–Sidak’s multiple comparison test. (I, J) *In vitro* spinogenesis of wild-type or *Inka2^-/-^* PCNs. Embryonic PCNs prepared from wild-type or *Inka2^-/-^* were electroporated with EGFP to visualize the dendritic spines. Synapse maturation is monitored at 7 div (I) and 13 div (J) by immunostaining with anti-PSD95, a marker for the post-synaptic density. Insets are the higher views of differentiating dendrites, showing the emergence of PSD95^+^ synapses sites. (K) The number of PSD95^+^ spines at 13 div (per 30 μm length of neurite). ***P < 0.001; Welch’s t-tests. Scale bars, 500 μm in (A), 2 μm in (F), 20 μm in (B), (E), and (I), and 5 μm in (J).

Importantly, Nissl staining showed that the thick apical dendrites appeared lost or invisible in a considerable number of layer V pyramidal neurons of *Inka2*^-/-^ neocortices (Figure 6B, *arrows*). To determine the morphological defects of pyramidal neurons, we performed Golgi silver impregnation staining using six-month-old *Inka2*^-/-^ and littermate mice. In wild-type mice brains, each apical dendrite emerged from the apex of a pyramidal neuron and extended perpendicularly toward the upper layers as a thick process (Figure 6E). However, *Inka2*^-/-^ pyramidal cells in layer V often had abnormal apical dendrites with thinner, irregularly distorted, and wavy morphology (Figure 6E, *arrows*). In addition to apical dendrites, such abnormal bending was frequently observed at many dendrite arbors formed by apical dendrites (Figure 6E, *arrows*) and basal dendrites (Figure 6F, *arrows*). Moreover, closer inspection manifested that the number of dendritic spines strikingly decreased in six-month-old *Inka2*^-/-^ mice (number of spines/10 μm apical dendrite: wild-type, 16.2 ± 0.6; *Inka2*^-/-^, 11.8 ± 0.6; number of spines/10 μm basal dendrite: wild-type, 10.6 ± 0.6; *Inka2*^-/-^, 7.0 ± 0.5) (Figure 6F–H). In general, most spines formed on the dendritic arborization have a globular-shaped head (head) and a thin neck that interconnects the head to the shaft of the dendrite. However, the remaining dendritic spines in *Inka2*-/-neurons had smaller and irregularly distorted heads (Figure 6F). Decreased density of dendritic spines was observed in juvenile mice (one-month-old), implying the defect in the onset of spine formation (spinogenesis) rather than the degeneration of spines (number of spines per 10 μm of apical dendrite: wild-type, 13.7 ± 0.5; *Inka2*^-/-^, 9.3 ± 0.4; spines/10 μm of basal dendrite: wild-type,10.8 ± 0.4; *Inka2*^-/-^, 6.8 ± 0.4) (Figure 6F–H). To exclude the possibility that the effects of the spine abnormalities in *Inka2*^-/-^ neurons were secondary to neuronal degeneration, we measured the cell density in cortical layer V of 12-month-old mice. The results showed no significant decrease or loss of neurons in the *Inka2*^-/-^ cerebral cortex (Figure S5A).

To confirm the impaired spinogenesis of *Inka2*^-/-^ neurons, the primary cortical neurons (PCNs) were prepared from the embryonic cortices and cultured *in vitro*. EGFP expression vector was introduced into the dissociated neurons from *Inka2*^-/-^ or *Inka2*^+/+^ embryos to visualize the fine morphology of each neuron. At 3 days *in vitro* (div), we could not observe any morphological difference between *Inka2*^-/-^ and wild-type neurons, which exhibited the extension of SMI312^+^ axon and normal arborization of MAP2^+^ dendrites (Figure S4). As early as 7 div, when active synaptogenesis and post-synaptic density protein 95 (PSD95) expression emerged in spines as a granular signal in wild-type cortical neurons, PSD95^+^ speckles remained inevident on *Inka2*^-/-^ neurons (Figure 6I). Instead, abnormally diffuse expression of PSD95 was observed within the *Inka2*^-/-^ dendritic shafts. At 13 div, when most neurons had fully differentiated and made synaptic contacts with one another, wild-type neurons formed a lot of EGFP^+^ dendritic spines that were colocalized with PSD95, which is a post-synaptic scaffolding protein used as a synaptic marker of excitatory neurons (Figure 6J). In contrast, *Inka2*^-/-^ neurons formed fewer PSD95^+^ spines (number of spines per 30 μm of dendritic shaft: wild-type, 12.5 ± 0.3; *Inka2*^-/-^, 8.9 ± 0.3) (Figure 6J, K). Altogether, these results indicated a critical function of Inka2 on dendritic spine formation.

### Inka2 represses LIMK–Cofilin signaling pathway in neurons

Our qPCR analysis showed that *Inka2* mRNA was considerably expressed in cultured mouse embryonic fibroblasts (MEFs) (Figure 7A) as well as the PCNs and the cerebral cortex of wild-type mice (Figure 7B) and that its expression was completely eliminated in *Inka2*^-/-^ MEFs and *Inka2*^-/-^ cerebral cortex. Meanwhile, a high level of *Inka1* expression was observed only in MEFs. This *Inka1* expression level was unchanged between wild-type and *Inka2*^-/-^ MEFs. Both the cerebral cortex and the cultured PCNs exhibited no expression of *Inka1* (Figure 7B). These findings suggested redundancy between the two genes in MEFs, while *Inka2* solely functions in the neural tissues.

We hypothesized that altered downstream pathways of Pak signaling contributed to the spinogenesis defects observed in the *Inka2*^-/-^ brain. LIMK–cofilin signaling is a major downstream pathway of Pak; Pak4 mediates the phosphorylation and activation of LIMK (pLIMK), leading to the phosphorylation and inactivation of cofilin (pCofilin) to promote stabilization and polymerization of the actin cytoskeleton (Ghosh et al., 2004). We first examined the pLIMK and pCofilin levels in MEFs prepared from subcutaneous tissue of wild-type or *Inka2*^-/-^ embryos. As a result, *Inka2*^-/-^ MEFs showed no significant change in pLIMK or pCofilin levels compared with wild-type MEFs (Figure 7C–E). Considering the possibility that Inka1 and Inka2 mutually suppress Pak4 signaling, we further performed the knockdown experiment of the *Inka1* gene in wild-type or *Inka2*^-/-^ MEFs using *Inka1* shRNAs (Figure S3B). The double knockdown of *Inka1* and *Inka2* significantly raised the content of pCofilin, despite the unchanged level of pLIMK (Figure 7C). Considering the redundancy of the two genes in MEFs, these results suggested functional cooperation of Inka1 and Inka2 in cofilin inactivation and actin dynamics in MEFs, mediated by the LIMK independent pathway.

Next, we investigated the phosphorylation levels of LIMK and cofilin in PCNs prepared from *Inka2*^-/-^ embryonic cortices. Both pLIMK and pCofilin levels were drastically upregulated in *Inka2*^-/-^ neurons at 7 div (Figure 7F–H). Contrary to the MEF case, *Inka1* knockdown in PCN showed no additive effect on phosphorylation of LIMK or cofilin (Figure 7F–H). This result might imply that Inka1 does not function in neurons, unlike fibroblasts. During the differentiation of cortical neurons, Inka2, but not Inka1, probably modulates the Pak4–LIMK–Cofilin pathway to regulate dendritic spine formation. In non-neuronal cells, such as fibroblasts, the Pak4 pathway might be cooperatively regulated by Inka1 and Inka2.

**Figure 7.**
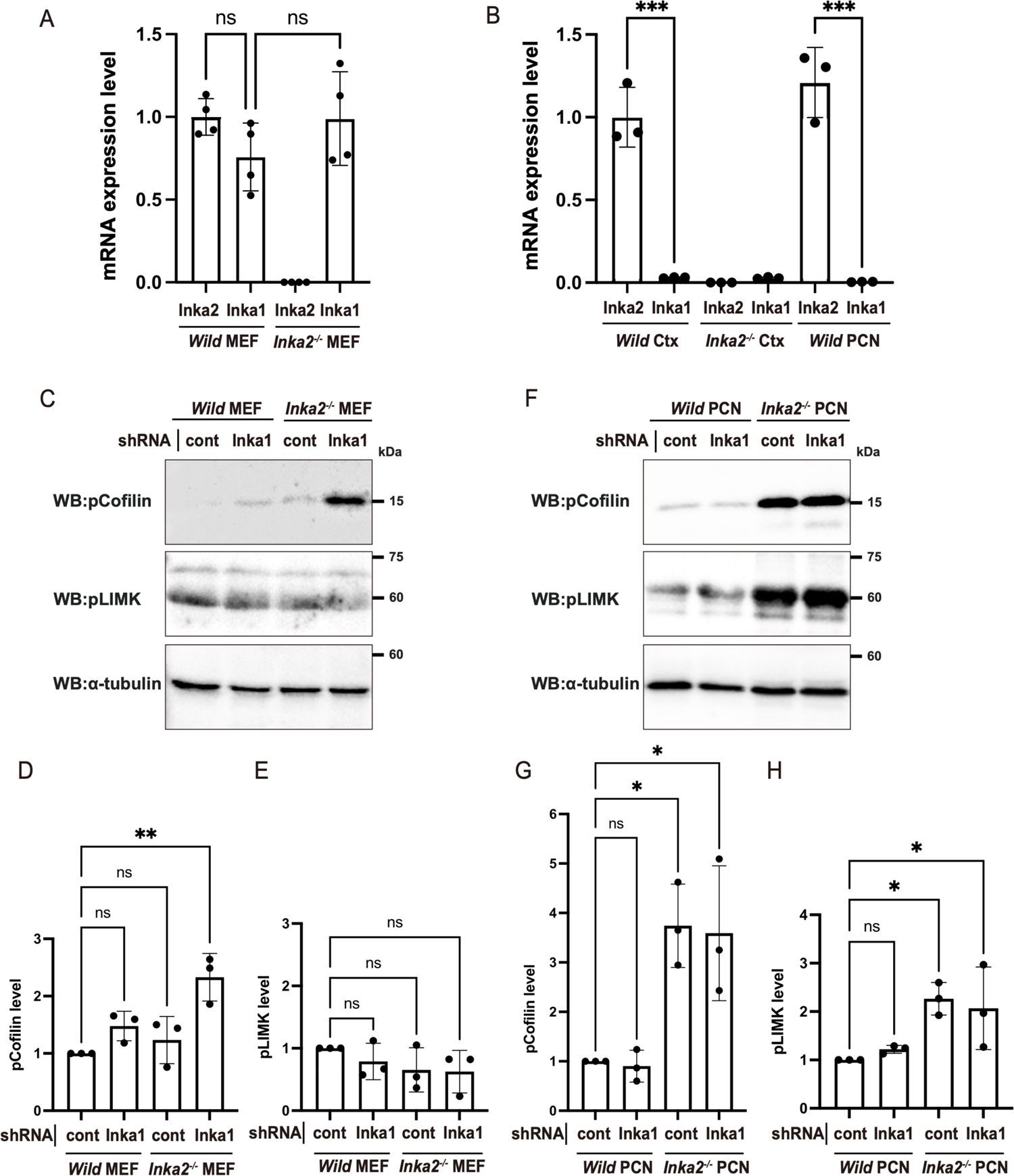
Upregulation of LIMK-Cofilin signaling in *Inka2^-/-^* neurons. (A, B) The expression level of *Inka2* and *Inka1* mRNAs in wild-type and *Inka2*^-/-^ MEFs (A), adult cerebral cortex (Ctx), and PCNs (B). Dots represent three independent qRT-PCR experiments normalized to the corresponding *β-actin* mRNA. ns, not significant; ***, P < 0.001; (A) Weltch’s t-test with Holm–Bonferroni correction. (B) One-way ANOVA. Holm–Sidak’s multiple comparisons test. (C–E) Knockdown of *Inka1* expression in *Inka2*^-/-^ MEF cells. The non-targeting control or *Inka1* shRNA was introduced into wild-type or *Inka2*^-/-^ MEF cells, and the phosphorylated forms of LIMK and cofilin were quantified by immunoblotting with anti-pLIMK, anti-pCofilin antibodies. Equal loading of lysate was verified by the immunoblot with an anti-α-tubulin antibody. (D, E) Quantification of the pLIMK (E) and pCofilin (D) levels normalized by α-tubulin. ns, not significant; **, P < 0.01; One-way ANOVA. Holm–Sidak’s multiple comparisons test. (F–H) Knockdown of *Inka1* expression in *Inka2*^-/-^ PCNs. The non-targeting control or *Inka1* shRNA was electroporated into the PCNs prepared from wild-type or *Inka2*^-/-^ embryonic cortices. At 7 div, each cell lysate was subjected to immunoblotting with anti-pLIMK, anti-pCofilin, and anti-α-tubulin antibodies. (G, H) Quantified comparison of the pLIMK (H) and pCofilin (G) level normalized by α-tubulin. ns, not significant; *, P < 0.05; One-way ANOVA. Holm–Sidak’s multiple comparisons test.

To confirm the *in vivo* effect of *Inka2^-/-^* in LIMK–Cofilin signaling in synapses, we isolated the synaptoneurosome (SN) fraction, in which pre- and post-synaptic sites are enriched (Figure 8A). Concentrated PSD95 protein level indicated that SNs were accurately fractionated from the cerebral cortices of *Inka2^-/-^* or wild-type mice (Figure 8B). Both pLIMK and pCofilin levels were drastically elevated in *Inka2*^-/-^ SN (Figure 8B–D). We further fractionated F and G actin from SN. Nonetheless, we could not detect a drastic change in the F/G actin ratio (data not shown) in *Inka2*^-/-^ SN. Altogether, Inka2 acts as a neuron-specific inhibitor of the Pak4 signaling cascade. During the differentiation of cortical neurons, Inka2, but not Inka1, modulates the Pak4–LIMK– Cofilin pathway to regulate neuronal development, including dendritic spine formation (Figure 8E).

**Figure 8.**
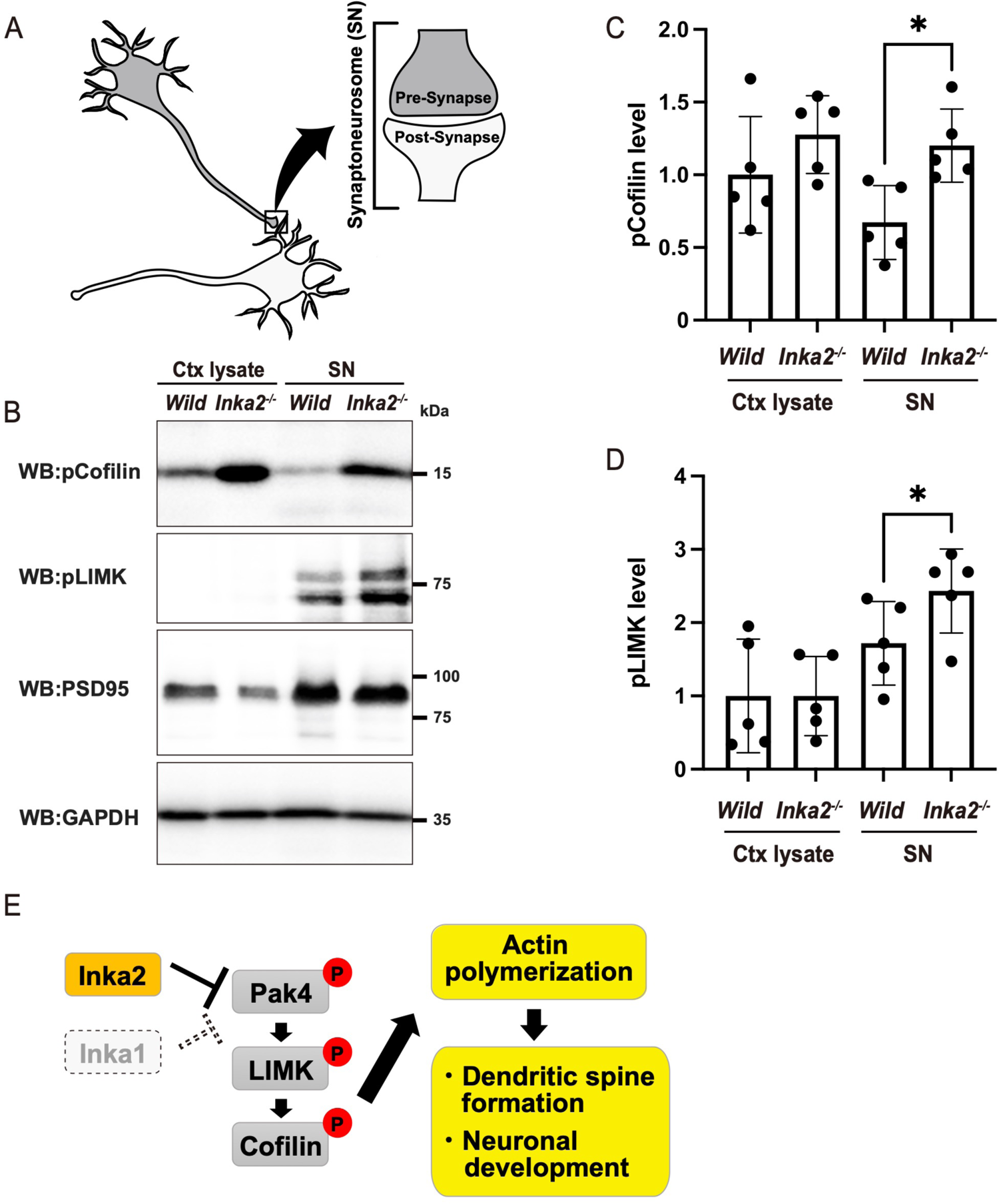
*Inka2* depletion in the brain enhances the LIMK-Cofilin signaling cascade in synaptoneurosome. (A) Schematic model of synaptoneurosome (SN). (B–D) Phosphorylation of LIMK and Cofilin were elevated in *Inka2*^-/-^ SN. Total protein lysate of the cerebral cortex (Ctx lysate) and the SN fraction were prepared from wild-type or *Inka2*^-/-^ and subjected to immunoblotting with anti-pCofilin, pLIMK, PSD95, and GAPDH antibodies. (C, D) Quantified comparison of pCofilin (C) and pLIMK (D) level normalized by GAPDH content. Five independent experiments were performed. *, P < 0.05; Paired t-test. (E) Model for the Inka2 function in cortical neurons. Inka2, but not Inka1, inhibits Pak4 activity to suppress the LIMK/Cofilin pathway in neurons affecting dendritic spine formation and neuronal development via actin dynamics.

### Translational repression of *Inka2* mRNA through G-quadruplex and GU-rich sequences

Northern blot and *in situ* hybridization analysis revealed the abundant expression of *Inka2* mRNA in the nervous system, especially in adult forebrain neurons (Iwasaki et al., 2015). Consistently, the present study showed the robust expression of lacZ reporter in forebrain neurons in *Inka2*^flox/+^ mice (Figure 5). A search on the integrated proteome databases (Global Proteome Machine Database, https://www.thegpm.org/) confirmed the presence of Inka2 peptides with the coverage of 84% (as derivatives from mouse Inka2; ENSMUSP00000095874) in mouse brain and neurons, indicating the occurrence of Inka2 protein synthesis in mouse brain. Nonetheless, as mentioned above, detecting the endogenous Inka2 protein using the antibodies in nervous tissues remains challenging.

This challenge may occur from the translational repression machinery of *Inka2* mRNA, allowing an undetectable expression level of endogenous Inka2 protein under the physiological condition. We found three G-quadruplexes (G4) motifs and a GU-rich region containing two tandemly expanded GU-repeats (GU)_19–24_ in the open reading frame (ORF) and 3′-untranslated region (3′-UTR) of *Inka2* mRNA (Figure 9C). The importance of the G4 and GU-repeat sequences was further underscored by their strict conservation between mouse and human *Inka2* mRNAs. G4s are four-stranded structures formed in guanine-rich DNA and RNA sequences (G_3_N_1–7_G_3_N_1–7_G_3_N_1–7_G_3_, N is any nucleotide base, including guanine). Because of square planar G-quartet assembly through Hoogsteen base pairing (not Watson–Crick base pairing) and stabilization by a central potassium ion, G4 is a globularly folded and thus an extremely stable structure. Recent accumulating studies have indicated that the G4 motif in 3′-UTR and ORF represses the mRNA translation in most cases. Several RNA binding proteins (RBPs), such as FMR1, DEAH-box helicase 36 (DHX36), Fused in sarcoma (FUS), and Aven, bind G4s to unwind G4 RNA and promote mRNA translation. In contrast, transactive response DNA-binding protein of 43 kDa (TDP-43), which is a candidate gene responsible for diverse neurodegenerative diseases, including amyotrophic lateral sclerosis (ALS) and frontotemporal lobar degeneration (FTLD), preferentially binds RNAs via a GU dinucleotide repeat (GU)_n_ element and regulates mRNA splicing, stability, and translational repression. As several RBPs, including FUS, FMR1, and TDP-43, are involved in synaptic functions by regulating the translation of target mRNAs (Davis and Broadie, 2017), we hypothesized the translational repression of *Inka2* mRNA by these RBPs. To test this idea, RNA-immunoprecipitation (RIP) experiment was performed. We demonstrated that *Inka2* mRNA was specifically immunoprecipitated with anti-FMR1 and anti-TDP-43 antibodies from the lysate of N2a cells (Figure 9A) and the mouse cerebral cortices (Figure 9B), indicating the *in vivo* binding of *Inka2* mRNA with the ribonucleoprotein (RNP) complex containing FMR1 and TDP-43. Next, we performed an RNA pull-down (RiboTrap) assay to isolate the RBPs associated with *Inka2* mRNA in brain extract. After incubating 5-Bromo-UTP (BrU)-incorporated *Inka2* mRNA with a cytoplasmic extract from N2a cells, BrU-*Inka2* RNA-RBP complexes were immunoaffinity purified. Then, the RBPs were identified by immunoblotting. Figure 9A and 9B show the association of *Inka2* mRNA with FMR1 and TDP-43. To determine the RNA elements required for the interaction with FMR1 and TDP-43, various *Inka2* RNA fragments (F1–F6) were synthesized and used for the RNA pull-down assay using N2a cell lysate. As shown in Figure 9C, FMR1 interacted with the RNAs F3, F4, and F5, which commonly contained two G4s. As RNA fragment F6 has little binding competence, FMR1 likely binds the tandem repeat of G4 elements rather than a single G4 motif or GU stretch in 3′-UTR (Figure 9C). In contrast, TDP-43 robustly interacted with RNAs F3, F5, and F6, indicating that TDP-43 specifically binds (GU)_19-24_ repeats in *Inka2* 3′-UTR (Figure 9C).

**Figure 9.**
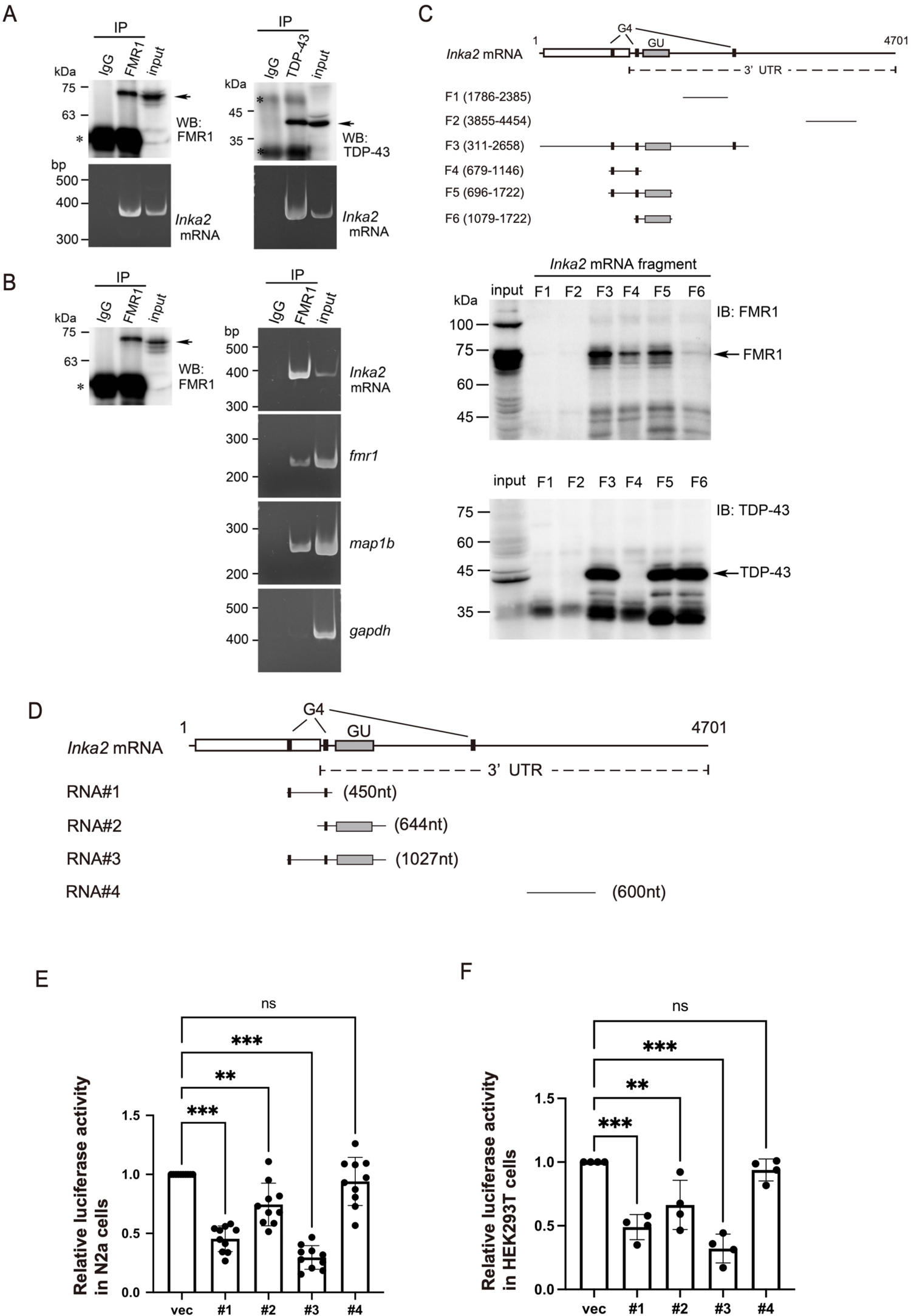
Translational repression of *Inka2* mRNA by RBPs. (A, B) RNA-immunoprecipitation (RIP) analysis showing the interaction between *Inka2* mRNA and the RNA-binding proteins FMR1 and TDP-43. Cell lysate from N2a cells (A) and adult mouse cerebral cortex (B) were subjected to immunoprecipitation (IP) with an anti-FMR1 or anti-TDP-43 antibodies to purify the ribonucleoprotein complex containing FMR1 and TDP-43. *Inka2* mRNA was detected by RT-PCR. The lower panels in A and the right panels in B represent the RT-PCR products separated by agarose gel electrophoresis. As FMR1 binds *map1b* mRNA, the *map1b* and *gapdh* mRNAs were used as a positive and negative control, respectively. The upper panels in A and the left panels in B show the immunoblot of IP and input lysate. (C) RNA pull-down assay using the *Inka2* mRNA fragments containing G4 or GU-repeat element. The schematic diagram represents the three G4s (filled square) and GU-repeat element (gray rectangle) in *Inka2* mRNA. An open square indicates the *Inka2* ORF. The indicated *Inka2* cDNA fragments (F1–F6) were *in vitro* transcribed for RNA pull-down assay. Each BrU-labeled *Inka2* RNA was incubated with N2a cell lysate, followed by IP with anti-BrdU antibody to precipitate the RNA fragment‒protein complexes. Immunoprecipitated proteins were analyzed by immunoblotting with anti-FMR1 or anti-TDP-43 antibodies (lower panels). (D–F) Luciferase reporter assay, showing the translational repression by the *Inka2* mRNA elements. (D) Depicted regions of *Inka2* mRNA (#1‒#4) were inserted downstream of the firefly luciferase gene. (E, F) Each dual-luciferase construct pmirGLO-*Inka2* mRNAs (#1–#4) was transfected to N2a (E) or HEK293T (F) cells, and the luciferase activity was measured. Firefly luciferase activity was normalized by the hRenilla luciferase activity and represented as the relative luminescent units (RLU). Five (for HEK293T) and 10 (for N2a) independent experiments were performed. ns, not significant; **, P < 0.01; ***, P < 0.001; One-way ANOVA. Holm–Sidak’s multiple comparisons test.

To examine whether the inhibition of Inka2 translation is dependent on G4s or GU elements, a dual-luciferase reporter assay was performed using HEK293T and N2a cells expressing various regions of *Inka2* mRNA downstream of the firefly luciferase gene (luc2) (Figure 9D). G4s and GU repeat elements efficiently suppressed the luciferase reporter translation in N2a (Figure 9E) and HEK293T cells (Figure 9F).

Altogether, these results demonstrate that Inka2 translation is inhibited by RBPs, including FMR1 and TDP-43, under physiological conditions. A strictly controlled *Inka2* level might be needed for neuronal differentiation and dendritic spine formation during normal brain development. We further performed knockdown experiments using the shRNAs specific to FMR1 and TDP-43 in N2a cells or primary cultured neurons but failed to detect a significant upregulation in the protein level of endogenous Inka2 (data not shown), implying the involvement of additional RBPs in translational regulation of Inka2.

## Discussion

### Inka2 acts as a neuronal inhibitor of Pak4 to regulate actin dynamics

In this study, we reported Inka2 as a novel and specific inhibitor of Pak4, which drives the depolymerization of F-actin through Pak4‒LIMK‒Cofilin signaling. The 38 amino-acid of the iBox sequence (Inka2-iBox) serves as an inhibitory domain of Inka2 and directly binds the catalytic domain of Pak4. *In vitro* analyses demonstrated that Inka2-iBox exhibits an IC_50_ of 60.3 nM, comparable to Inka1-iBox (67.7 nM) (Figure 4). A previous study has suggested that Inka1 specifically inhibits Pak4, not Pak1 (Baskaran et al., 2015). Similarly, Inka2 efficiently inhibits Pak4 activity but does not affect Pak1, implying that the Inka family is a specific inhibitor for group II Pak (Pak4-6).

Previous studies have demonstrated the central function of Pak4 signaling in regulating actin dynamics and cell shape. Overexpression of constitutively active Pak4 (S445N), LIMK1, or cofilin induces cytoskeletal changes, including rounded, lacked stress fibers, and contains actin clusters in the peripheral cellular regions (Dan et al., 2001; Qu et al., 2001). In contrast, constitutively active PAK1 (T423E) phosphorylated LIMK1 inefficiently compared with PAK4 (Dan et al., 2001). In this study, Inka2, but not Inka1, was shown to play a vital role in cell morphology dependent on Pak4 signaling. Forced expression of Inka2 caused cell-rounding of HEK293T, and this phenotype was partially rescued by Pak4 overexpression. Moreover, Inka2 overexpression inhibited the abnormal extension and formation of cellular processes induced by Pak4cat expression in HeLa cells. Considering the elevated phosphorylation of LIMK and cofilin in *Inka2*^-/-^ neurons, the effect of Inka2 on cell morphology is highly likely by regulating Pak4/LIMK/Cofilin signaling cascade. Nevertheless, Inka1 overexpression did not cause any morphological change in cells, despite the similar inhibitory effect of Inka2-iBox and Inka1-iBox on Pak4 kinase activity. This discrepancy may be partly due to the difference in subcellular localization of Inka1 and Inka2. Inka2 mainly colocalizes with F-actin in the cytosol; however, Inka1 is predominantly localized in the nucleus (Figure 1). Since Inka2ΔCC is localized primarily in the nucleus (Figure 1E), the N-terminal CC domain of Inka2 may be necessary for the cytoplasmic localization, enabling interaction with cytoplasmic Pak4. Considering both Inka1 and Inka2ΔCC translocate from the nucleus into the cytoplasm upon co-expression with Pak4 (Figure 2 F), there might be another molecular machinery that transports the Inka family proteins to cytosol dependent on the Pak4 expression.

Besides cell-shape regulation, Inka2-mediated actin dynamics is closely related to various cellular processes, including cell migration, proliferation, cancer invasion, and adherent junction formation. Inka2 is the direct target of tumor suppressor gene and transcription factor p53, and Inka2 expression decreases cancer cell growth by inhibiting the β-catenin signal (Liu et al., 2019). In humans, FAM212b (INKA2) expression level is correlated with increased melanoma metastasis and poor clinical outcomes in patients (Chen et al., 2021). We recently demonstrated that *Inka2* knockdown accelerates cell migration with increased focal adhesion turnover in NIH3T3 cells (Akiyama et al., 2020). Since such accelerated cell migration is also induced by LIMK1 and Pak4 overexpression (Ahmed et al., 2008), Inka2‒Pak4‒LIMK cascade controls the driving force of cell migration. Meanwhile, accumulated evidence suggests the crucial and comparable function of Pak4 in cell proliferation, adherence junction formation, and cell migration (Ahmed et al., 2008; Qu et al., 2003; Tian et al., 2011). Similar to the effect of Inka2 on tumor metastasis, Pak4 inhibitor PF-3758309 blocks the growth of multiple tumor xenografts (Murray et al., 2010). Altogether, our results strongly suggest the involvement of Inka2 in the Pak4 signaling pathway modulating the diverse cellular processes accompanied by the reorganization of F-actin.

### Inka2 regulates synapse formation

Transient overexpression of Rac1, Pak, and LIMK induces the formation of dendritic spines in cultured pyramidal neurons (Hayashi et al., 2007; Meng et al., 2002; Wiens et al., 2005; Zhang et al., 2005). A decrease in the dendritic spine density has been reported in the cerebral cortex of patients with schizophrenia. Disrupted-in-Schizophrenia-1 (DISC1), a candidate responsible gene for pathogenesis underlying schizophrenia, plays a role in the post-synaptic density associated with Rac1 (Hayashi-Takagi et al., 2010). The transient and short-term depletion of DISC1 activates Rac1 and induces spine growth in primary cultured neurons. In contrast, prolonged knockdown of DISC1 expression leads to a reduction in spine density in the cortical neurons, analogous to the synaptic pathology of schizophrenia (Hayashi-Takagi et al., 2010). Moreover, the chemical inhibitors for PAKs prevent the spinogenesis impairment induced by DISC1 deficiency *in vitro* and *in vivo* (Hayashi-Takagi et al., 2014). These observations imply that the constitutive activation of the Rac1–Pak pathway reduces dendritic spine density.

In mammals, Pak4 is expressed ubiquitously in various tissues, including the brain (Qu et al., 2003). Other members of group II Paks, Pak5 and Pak6, are also highly and specifically expressed in the CNS (Civiero and Greggio, 2018; Nekrasova et al., 2008). Given the similar substrate properties of Pak5 and Pak6 to Pak4, Inka2 may act as a common inhibitor of group II Paks. In contrast, *Inka1* expression is principally observed in the migratory neural crest cells, dorsal root ganglion, branchial arches, and mesenchyme cells of the developing brain, but not in the neural tissue of the CNS itself (Figure 5L, 7B) (Reid et al., 2010). Predominant *Inka2* expression in the CNS (Iwasaki et al., 2015) presumably reflects an equivalent or similar Inka1 and Inka2 function in different non-overlapping regions as Pak4 or group II Paks repressors. Our previous *in situ* hybridization analysis revealed that *Inka2 mRNA* expression emerges on pyramidal neurons at P7 mice cerebral cortex and becomes prominent in the forebrain neurons as adult stage (Iwasaki et al., 2015). Coincidently, our present X-gal staining and *in situ* hybridization showed specific and robust *Inka2* expression in the forebrain neurons, including pyramidal cells in the cerebral cortex. Histological analysis of *Inka2*^-/-^ brain showed abnormal dendrites bending and spinogenesis impairment in pyramidal neurons. Given that F-actin is a major cytoskeletal component of the dendritic spine and that local actin remodeling of dendrites alters spine plasticity (Borovac et al., 2018), postulating that the dysregulated Rac–group II Paks signaling and deviating actin dynamics cause the abnormal morphology and deterioration of dendritic spines in *Inka2*^-/-^ neurons are reasonable. *Inka2* deficiency, but not *Inka1*, led to the elevated activation of LIMK and cofilin signaling in cortical neurons *in vivo* and *in vitro*, allowing impaired spinogenesis. In contrast, enhanced cofilin activation was observed in MEF cells only when both *Inka1* and *Inka2* were knocked down (Figure 7). This cell-type-dependent activation of cofilin might support the notion that Inka1 and Inka2 cooperatively and redundantly regulate Pak4 signaling in non-neuronal cells.

*Pak4* KO mice resulted in embryonic lethality due to the defects in cardiovascular and nervous systems (Qu et al., 2003), indicating that the molecular machinery involved in the Pak4 signaling must be tightly controlled *in vivo*. We showed the direct association of *Inka2* mRNA with FMR1 and TDP-43, which functions as a translational repressor in neurons. FMR1 and TDP-43 bind G4 and GU-rich repeats, respectively, inhibiting the translation of synaptic proteins, such as PSD95, Map1b, Arc, CaMKII, and metabotropic glutamate receptors (Darnell et al., 2001; Davis and Broadie, 2017). Using the luciferase reporter assay, we demonstrated that G4s and GU-rich repeats in *Inka2* mRNA repress the protein translation *in vitro*. Based on these findings, we speculate that the translational repression of *Inka2* mRNA in neurons by the RBP network, including FMR1. FMR1 KO mouse, an FXS model, shows an increased number of immature dendritic spines, resulting in learning and memory deficits. Pak signaling and F-actin stability were enhanced in the dendritic spines of FMR1 KO mice (Chen et al., 2010). The increased translation of FMR1 target mRNAs in the dendritic spine caused by FMR1 deficiency is presumed to be the source of pathology underlying FXS (Davis and Broadie, 2017). As a target mRNA of FMR1, *Inka2* may be involved in the decreased Pak signaling and aberrant spine formation in FXS patients. However, *Inka2* mRNA is possibly associated with additional RBPs, such as TDP-43, FUS, and DHX36. To determine whether Inka2 translation is concomitantly controlled by combining other RBPs implicated in different neuropsychiatric disorders is intriguing.

## Materials and Methods

### Animals

ICR and C57BL/6J mice purchased from Japan SLC, Inc (Shizuoka, Japan) were housed under temperature- and humidity-controlled conditions on a 12/12 h light/dark cycle, with *ad libitum* access to food and water. The date of conception was established using a vaginal plug and recorded as embryonic day zero (E0.5). The day of birth was designated as P0.

Chimera mice with *Inka2*^tm1a^ were created using blastocyst injection of targeted ES cell clone (EPD0805_2_E10) from KOMP (www.komp.org). The KO-first allele, designated as tm1a, is a preconditional null (non-expressive) allele containing a lacZ reporter cassette with an internal ribosome entry site (IRES) and a simian virus 40 polyadenylation site, allowing the transcriptional termination at the upstream of critical exon (Skarnes et al., 2011). The critical coding exon of *Inka2* (exon 2) and the neo selection cassette are flanked by loxP sites (floxed) in the *Inka2*^tm1a^ allele. In heterozygous mice carrying the tm1a allele, lacZ activity serves as a reporter for endogenous expression of each targeted gene by splicing the cDNA to the lacZ cassette. We further converted the tm1a allele to a KO reporter allele (*Inka2*^tm1b^) by crossing *Inka2*^tm1a^ heterozygous mice with CAG-Cre driver mice (B6.Cg-Tg[CAG-Cre]CZ-MO2Osb, RBRC01828; RIKEN BRC, Tokyo, Japan) harboring the transgene of CAG-promoter-driven Cre recombinase (Matsumura et al., 2004). *Inka2*^tm1b^ is the lacZ-tagged KO (null) allele, in which the critical exon (exon 2) was eliminated by Cre/loxP-mediated excision. For brevity, homozygous alleles of *Inka2*^tm1a^ and *Inka2*^tm1b^ are hereafter referred to as *Inka2*^flox/flox^ and *Inka2*^-/-^. After eight to nine generations of backcrossing with C57BL/6J, the progeny of heterozygous intercrosses was used for analyses. All protocols were approved by the Committee on the Ethics of Animal Experiments of Waseda University.

### X-gal-staining

Wild and *Inka2*^flox/+^ mice were transcardially perfused with saline solution, followed by 2% paraformaldehyde (PFA) and 0.2% glutaraldehyde in 0.1 M phosphate buffer (pH 7.4). Free-floating frozen sections of adult brains were cut at 30 μm thickness using a cryostat. The sections were washed with phosphate-buffered saline (PBS) and then twice with X-gal wash buffer (PBS, 2 mM MgCl_2_, 0.01% sodium deoxycholate, and 0.1% NP-40) for 10 min at 4℃. X-gal staining was performed in the X-gal wash buffer containing 5 mM potassium ferricyanide, 5 mM potassium ferrocyanide, and 1 mg/mL X-gal for 2–24 h at 37℃. Free-floating sections were perfused with 4% PFA for 15 min and collected on MAS coated glass slides (Matsunami Glass, Osaka, Japan). Sections were counterstained with neutral red, dehydrated, and coverslipped with Entellan new (Merck Millipore, Burlington, MA, USA).

### In situ hybridization

*In situ* hybridization was performed as described previously (Iwasaki et al., 2015). The cDNA fragments corresponding to 495–1234 nucleotide (nt) of mouse *Inka2* cDNA (GenBank accession no. BC052371) and 170–1015 nt of mouse *Inka1* cDNA (accession no. NM_026597) were subcloned and used to prepare the DIG-labeled antisense riboprobes. Hybridization signals were detected using alkaline phosphatase-conjugated anti-DIG antibody (1:1000, Merk Millipore), followed by incubation with 0.11 mM nitro blue tetrazolium and 0.12 mM 5-Bromo-4-chloro-3-indoyl phosphate in 6% polyvinyl alcohol, 100 mM NaCl, 100 mM Tris–HCl, pH 9.8 and 50 mM MgCl_2_ at 37 ℃. After color development, the sections were dehydrated using ethanol and xylene and then cover-slipped with Entellan New (Merck Millipore).

### Nissl staining

*Inka2*^-/-^ mice and the wild-type littermates were perfused transcardially with saline solution, followed by 4% PFA in 0.1 M phosphate buffer (pH 7.4). Adult mice brains were cut at 30 μm thickness and collected on MAS coated glass slides. After washing the sections with distilled water, the sections were degreased using 100% ethanol (EtOH) for 10 min. The sections were again washed with water and reacted with 0.1% Cresyl Violet (Merck Millipore) for 10 min at 25°C. Sections were then decolorized using 70% EtOH, dehydrated, and coverslipped with Entellan New.

### Golgi staining

Golgi silver staining was performed using the FD Rapid GolgiStain Kit (FD Neuro Technologies, Columbia, MD, USA) according to the manufacturer’s protocol. Images were acquired using a DP72 CCD camera (Olympus Corporation, Tokyo, Japan) mounted on an ECLIPSE E800 microscope (Nikon, Tokyo, Japan).

### Plasmids

cDNA fragments corresponding to *Inka1* and *Inka2* ORFs were isolated using RT-PCR of the RNAs of E12 embryonic and adult mice brains and subcloned in-frame into a pEGFP-C2 expression vector (Takara Bio Inc., Shiga, Japan). To construct the CAG promoter-driven *Inka2* in mammalian cells, the EGFP-Inka2 ORF fragment was subcloned into the pCAG-GS vector (Yamada et al., 2020). Deletion mutant EGFP-Inka2ΔiBox and EGFP-Inka2ΔCC were constructed from pEGFP-Inka2 using Q5 Site-Directed Mutagenesis Kit (New England BioLabs, Ipswich, MA, USA). Expression plasmids for Pak1 (pCMV6M-Pak1) and Pak4 (pWZL-Neo-Myr-Flag-Pak4) were obtained from Addgene (plasmid #12209 and # 20460, respectively; Watertown, MA, USA) (Sells et al., 1997). For Pros2-His-Pak1cat expression in E. coli, Pak1 fragment (amino acids 270–521) was PCR-amplified from pCMV6M-Pak1 and subcloned into pCold-Pros2 vector (Takara Bio Inc.) to generate pCold-Pros2-Pak1cat. pXJ40-Flag-PAK4, pXJ40-Flag-PAK4cat (amino acids 286–591), pSY5M-Pak4cat (286–591), pGEX-Raf13, pGEX-4T1-Inka1-iBox-38 (166–203), and pGEX-4T1-Inka2-iBox-61 (133–193) (Baskaran et al., 2015) were provided by Dr. E. Manser and Dr. Y. Baskaran (Institute of Molecular and Cell Biology, Singapore). We purchased a MISSION shRNA vector library encoding the microRNA-adapted shRNA targeting mouse *Inka1* (Sigma-Aldrich, St. Louis, MO, USA). Among five shRNA clones (TRCN0000269004, TRCN0000269005, TRCN0000283867, TRCN0000283870, and TRCN0000202275), TRCN0000269004 and TRCN0000202275 were designated as shRNA #1 and shRNA #2, respectively. The targeting sequences of shRNA #1 and shRNA #2 were: 5′-GTAGCAGAAGACAAGGGTTTA-3′ and 5′-GCGTTTCTCAGTCAGTGACAT-3′, respectively.

### Primary antibodies

anti-GFP (chicken polyclonal IgY, GFP-1010, Aves Labs, Tigard, OR, USA; 1:2000 for ICC), anti-GFP (rabbit polyclonal, GTX113617, GeneTex, Irvine, CA, USA; 1:2000 for WB), anti-GFP (rabbit polyclonal, NB600-308, Novus Biologicals, Littleton, CO, USA; 1:500 for IP), anti-DDDDK (Flag) (mouse monoclonal, FLA-1, MBL Life Sciences, Tokyo, Japan; 1:10000 for ICC and WB), anti-β-actin (mouse monoclonal, 2D4H5, Proteintech, Rosemont, IL, USA; 1:10000 for WB), anti-α-tubulin (rabbit polyclonal, #11224-1-AP, Proteintech; 1:5000 for WB), anti-TDP-43 (rabbit polyclonal, RN107PW, MBL; 1:1000 for WB), anti-FMP1 (rabbit polyclonal, RN016P, MBL; 1:1000 for WB), anti-PSD95 (mouse monoclonal, 6G6-1C9, Abcam, Cambridge, UK; 1:500 for ICC, 1:3000 for WB), anti-pCofilin (Ser3) (rabbit monoclonal, 77G2, Cell Signaling Technology, Danvers, MA, USA; 1:1000 for WB), anti-pLIMK1 (Thr508)/LIMK2 (Thr505) (rabbit polyclonal, #3841, Cell Signaling Technology; 1:1000 for WB), anti-MAP2(chicken polyclonal IgY, Poly28225, Biolegend, San Diego, CA, USA; 1:10,000 for ICC), and anti-SMI312 (mouse monoclonal, 837904, Biolegend; 1:1000, for ICC).

### Cell culture

HEK293T, N2a, HeLa, and MEF cells were cultured in Dulbecco’s Modified Eagle Medium (DMEM) containing 10% fetal bovine serum (FBS), penicillin/streptomycin, and L-glutamine. For the primary cortical neurons (PCNs), embryonic cerebral cortices from E16.5 were dissected, as previously described (Yamada et al., 2020). Cells were seeded onto poly-D-lysine-coated dishes (Sigma-Aldrich) and cultured in a neurobasal medium containing 2% B27 (Life Technologies, Carlsbad, CA, USA) and 1% GlutaMax (Life Technologies) for 3‒13 div. For the primary culture of MEF cells, E12.5 embryos, excluding the head and internal organs, were dissociated and seeded in DMEM containing 10% FBS, penicillin/streptomycin, and L-glutamine.

### Cell transfection

Cell transfection was performed as previously described (Yamada et al., 2020). Cultured cells were transfected with plasmid DNA and PEI MAX (Polysciences, Warrington, PA, USA) complexes (ratio of DNA to PEI MAX, 1:3, w/w) formed in Opti-MEM I by incubating for 15 min at 25°C. The DNA complexes were added to cell cultures together with Opti-MEM for 3 h, followed by cultivation with serum-containing DMEM. PCNs were electroporated with plasmid DNA using a NEPA21 Electroporator (Nepagene, Chiba, Japan) according to the manufacturer’s protocols (two pulses of 275 V for 0.5 ms with an interval of 50 ms).

### In utero electroporation

In utero electroporation was performed as previously described (Yamada et al., 2021; Yamada et al., 2020). Pregnant mice were anesthetized *via* intraperitoneally injecting a mixture containing medetomidine, midazolam, and butorphanol. 5 μg/μL DNA solution in PBS with 0.01% Fast Green dye (Sigma-Aldrich) was injected into the lateral ventricle through the uterus wall, followed by electroporation. The following constructs were electroporated: pCAG-EGFP and pCAG-EGFP-Inka2. Electric pulses were generated using NEPA21 (Nepagene) and applied to the cerebral wall using a platinum oval electrode (CUY650P5, Nepagene), with four pulses of 35 V for 50 ms with an interval of 950 ms. An anionic electrode was placed on the lateral cortex to ensure the incorporation of DNA into the ventricular/subventricular zone (VZ/SVZ). Embryos were perfused at E15.5 with 4% PFA through cardiac perfusion.

### Immunostaining, actin staining, and TUNEL staining

Immunostaining was performed as previously described (Yamada and Sakakibara, 2018). Fixed cells were blocked for 1 h with 5% normal goat or donkey serum in PBST (0.0.5% Triton X-100 in PBS), followed by the incubation with primary antibodies in a blocking buffer for 1 h. After washing with PBST six times, sections were incubated for 1 h with Alexa Fluor 488-, Alexa Fluor 555-, Alexa Fluor 647-(Thermo Fisher Scientific, Waltham, MA, USA), or DyLight 488-, DyLight 549-(Jackson ImmunoResearch, PA, USA) conjugated secondary antibodies. After counterstaining with 0.7 nM Hoechst 33342 (Thermo Fisher Scientific), for actin staining, F-actin was stained using Acti-stain 555 phalloidin (Cytoskeleton, Inc., Denver, CO, USA), according to the manufacturer’s instructions. For detecting apoptosis, TUNEL staining was performed using the In Situ Cell Death Detection Kit, TMR-red (Roche Diagnostics, Basel, Switzerland), according to the manufacturer’s instruction. Images were acquired using a confocal (FV3000, Olympus) or fluorescence inverted microscope (AxioObserver, Zeiss, Germany).

### Immunoprecipitation (IP) and western blotting

IP and western blotting were performed as previously described (Yamada et al., 2021). For IP, cells were washed in ice-cold PBS and lysed in ice-cold lysis buffer containing 50 mM Tris-HCl (pH 7.5), 150 mM NaCl, 2 mM EDTA, 1% NP-40, and protease inhibitors (cOmplete Mini Protease Inhibitor Cocktail, Roche Diagnostics) for 30 min at 4°C. Lysates were centrifuged at 15,000 rpm for 10 min at 4°C, and the supernatants were coupled with protein A/G plus agarose IP beads (Santa Cruz Biotechnology, Dallas, TX, USA) overnight at 4°C. Rabbit or mouse IgG (Thermo Fisher Scientific) was used as a control. After brief centrifugation, beads were washed four times with 0.1% NP40 in PBS, and then the bound proteins were dissolved by treatment with 2×sample buffer (125 mM Tris-HCl, pH 6.8, 4% SDS, 10% sucrose, and 0.01% bromophenol blue). IP samples were resolved on 8%–12% SDS-PAGE and electroblotted onto Immobilon-P membranes (Merck Millipore) using a semidry transfer apparatus. After blocking with 5% skim milk in TBST (150 mM NaCl, 10 mM Tris– HCl, pH 7.4, 0.1% Tween 20), membranes were incubated with primary antibody for 1 h, followed by incubation with the horseradish peroxidase (HRP)-conjugated secondary antibody (Cytiva, Tokyo, Japan or Jackson ImmunoResearch). The signal was detected using the Immobilon Western chemiluminescent HRP substrate (Merck Millipore) and visualized using Fusion Solo S (Vilber Lourmat).

### Synaptoneurosome fraction preparation

Synaptoneurosome (SN) fraction was prepared as described (Villasana et al., 2006) with some modifications. Cerebral cortices of four-week-old mice were homogenized with SN buffer (10 mM HEPES pH 7.0, 1 mM EDTA, 2 mM EGTA, 0.5 mM DTT, and protease inhibitors), followed by filtration through a 70 μm-mesh cell strainer. The filtrate was further passed through a 5 μm pore-sized hydrophilic membrane using a Luer-lock syringe. After the filtrated sample was centrifuged at 1000 ×g for 10 min at 4°C, the pelleted SN fraction was dissolved in an SDS-sample buffer, followed by immunoblotting. Preparation of F/G actin from SN was performed as previously described (Pyronneau et al., 2017). Briefly, the SN fraction was resuspended in cold lysis buffer (50 mM PIPES pH 6.9, 50 mM KCl, 2 mM MgCl_2_, 1 mM EGTA, 0.2 mM DTT, 0.5% Triton X-100, 1 mM sucrose, and protease inhibitors) and centrifuged at 15,000 ×g for 30 min. F-actin and G-actin were recovered as the pellet and supernatant, respectively. Each fraction was dissolved in an equal volume of SDS-sample buffer, followed by immunoblotting.

### G-actin and F-actin fractionation

G-actin and F-actin fractions were obtained as described (Zhao et al., 2017) with some modifications. HEK293T cells expressing EGFP-Inka or Flag-Pak4 were lysed with actin stabilization buffer (50 mM PIPES at pH 6.9, 50 mM NaCl, 5 mM MgCl_2_, 5 mM EGTA, 5% glycerol, 0.1% NP40, 0.1% Triton X-100, 0.1% Tween 20, 0.1% β-mercaptoethanol, and 1 mM ATP) at 37°C for 10 min and centrifuged at 350 g for 5 min twice at 4°C. Supernatants were further ultracentrifuged at 100,000 g at 4°C for 1 h. The G-actin was recovered in the supernatant, and F-actin was enriched in a pellet. Each fraction was dissolved using an SDS-sample buffer, followed by immunoblotting. For *in vitro* cell-free reconstitution of F-actin, the HEK293T cell lysate was prepared with actin stabilization buffer and incubated with the purified proteins of GST or GST-Inka2-iBox (500 nM) together with His-Pak4cat (25 nM) for 30 min at 37°C. After incubation, the reaction mixture was centrifuged at 350 g for 5 min twice. The supernatants were further ultracentrifuged at 100,000 g for 1 h at 4°C to recover the G- and F-actin fractions.

### RNA-immunoprecipitation (RIP)

IP of endogenous RNAs was performed using the RIP-Assay Kit (MBL Life Sciences) according to the manufacturer’s instructions. In brief, cerebral cortices from adult C57BL/6J mice (0.2 g wet weight) or N2a cells (approximately 2 × 10^7^ cells) were washed with ice-cold PBS three times and lysed in 500 μL lysis buffer (+) supplemented with 1.5 mM dithiothreitol (DTT), 10 U/mL RNase inhibitor (Promega, Madison, WI, USA) and cOmplete Mini Protease Inhibitor Cocktail. After incubating for 10 min at 4℃ on ice, cell lysate was collected by centrifugation at 12,000 × g for 5 min at 4℃. The cell lysate was precleared by incubating Protein A/G Plus-agarose beads (sc-2003, Santa Cruz Biotechnology) for 1 h at 4℃. After centrifugation at 2,000 g for 5 min at 4°C, the supernatant was mixed with 25 μL (50% slurry) of Protein A/G Plus agarose beads conjugated to 15 μg of normal rabbit IgG (PM035, MBL), anti-FMR1 (RN016P, MBL), or anti-TDP43 (RN107PW, MBL) antibody. After incubation with rotation for 3 h at 4°C, each bead sample was washed with 1 mL wash buffer (+) containing 1.5 mM DTT four times and split 4/5 for RNA and 1/5 for protein fraction. For isolation of the protein fraction, beads were resuspended in 20 μL of 1× SDS PAGE sample buffer and boiled for 5 min for western blotting analysis. The RNA fraction was eluted from agarose beads and redissolved in 20 μL nuclease-free water according to the manufacturer’s instructions. After the RQ1 DNase (Promega) treatment, eluted RNAs were reverse-transcribed using PrimeScript™ 1st strand cDNA Synthesis Kit (Takara Bio) and subjected to PCR. Total RNA (input) prepared from cell or tissue lysate and the eluted RNAs from normal rabbit IgG controls were assayed simultaneously to verify the specificity of the detected signals (n = 4 for each experiment). Target sequences were amplified by 18–30 cycles of PCR using Ex-Taq™ (Takara Bio). For semiquantitative PCR, the number of amplification cycles was optimized to ensure that PCR products were quantified during the exponential phase of the amplification. A 10 μL aliquot of each PCR product was size-separated using electrophoresis on a 5% polyacrylamide gel (PAGE) with Tris-borate EDTA buffer. PCR primer sets used for detecting mouse *Inka2*, *FMR1*, *map1b*, and *gapdh* mRNA are given in Table S1.

### RNA pull-down assay

The RNA pull-down assay was performed using RiboTrap Kit (MBL), according to the manufacturer’s instructions. Briefly, the cDNA fragments corresponding to the coding region or 3′-UTR of mouse *Inka2* cDNA were subcloned into a pGEM-3Z plasmid vector (Promega) for *in vitro* transcription. The BrU‒labeled RNA was prepared using a CUGA 7 *in vitro* Transcription Kit (Nippon Gene, Japan). Purified BrU-labeled RNA (50 pmol each) was bound on the Protein G Plus-Agarose beads conjugated to an anti-BrdU monoclonal antibody. Cytoplasmic extract prepared from N2a cells (approximately 1 × 10^8^ cells) according to the manufacturer’s instruction (MBL) was precleared by the treatment with Protein G beads and then incubated with the BrU-labeled RNAs on antibody conjugated beads for 2 h at 4°C. After washing the sample beads four times with the low-ionic strength wash buffer I (+) supplemented with 1.5 mM DTT, BrU-RNA/protein complexes were eluted with 50 μL PBS containing BrdU by immunoblotting.

### Dual-luciferase reporter assay

To construct luciferase reporter plasmids for mouse *Inka2*, various cDNA fragments spanning the coding region and 3′-UTR were amplified by PCR and cloned individually downstream to the firefly luciferase 2 at the XbaI site of the pmirGLO Dual-Luciferase expression vector (Promega). For the luciferase reporter assay, 100 ng luciferase reporter plasmid was transfected into N2a or HEK293T cells cultured in 96-well plates using PEI MAX. At 36 h after transfection, cells were lysed with the Dual-Glo reagent (Promega), and firefly luciferase and hRenilla luciferase activities were determined using the Dual-Glo luciferase assay system (Promega) and Fluoroskan FL (Thermo) according to the manufacturer’s instructions. Firefly luciferase activity was normalized to hRenilla luciferase activity for each construct. For each transfection, luciferase activity was averaged from triplicates and the relative firefly luciferase activity was compared to that of the controls (empty pmirGLO transfected cells).

### 3′Rapid amplification of cDNA end (RACE)

Total RNA was isolated from 8-week-old wild-type or *Inka2^-/-^* mice forebrains using the RNAiso Plus reagents (Takara Bio). PolyA^+^ mRNA was extracted from total RNA using Oligotex TM-dT30 mRNA Purification Kit (Takara Bio), according to the manufacturer’s instructions. 3′RACE was performed using SMARTer RACE cDNA Amplification Kit (Takara Bio) per the manufacturer’s instructions. The cDNA fragment containing the *Inka2* exon1 sequence was amplified by PCR using *Inka2* gene-specific primer (GSP) and the nested primer (NGSP).

### Quantitative PCR

Quantitative PCR was performed as previously described (Yamada et al., 2021). Total RNAs prepared from adult mice cerebral cortex, PCNs at 10 div, and MEFs were reverse-transcribed using a PrimeScript II first-strand cDNA Synthesis Kit (Takara Bio) with a random hexamer primer. According to the manufacturer’s protocol, PCR was performed using the TB Green Ex Taq II Mix (Takara Bio) and the Thermal Cycler Dice Real-Time System (Takara Bio).

### Expression and purification of recombinant proteins

For the production of His_6_-Pak4cat proteins, pSY5M-Pak4cat were introduced into Escherichia coli strain BL21(DE3) pLysS (Promega), and the fusion proteins were induced by incubation with 0.5 mM isopropyl-b-D-thiogalactoside (IPTG) for 3.5 h at 30°C. pGEX-Raf13, pGEX-4T1-Inka1-iBox-38, and pGEX-4T1-Inka2-iBox-61 were used to produce GST-Raf13, GST-Inka1-iBox, and GST-Inka2-iBox, respectively. The recombinant His_6_-tagged-Pak1 catalytic domain was not expressed in the E. coli expression system using theT7 promoter-based expression vector, probably due to the toxicity to E. coli, as previously reported (Ng et al., 2010). Therefore, we employed the cold shock expression vector (pCold) (Tanabe et al., 1992), in which the recombinant protein is produced under the cold shock (cspA) promoter with Protein S (Pros2)-tag. The expression plasmid pCold-Pros2-Pak1cat was transformed into BL21(DE3) pLysS, and protein expression was induced using 0.5 mM IPTG for 20 h at 16°C. In this expression system, Pak1 catalytic domain (PH-Pak1cat) was successfully expressed and purified (Figure S3C). Recombinant proteins were affinity purified using Ni-Sepharose 6 Fast Flow (Cytiva) or Glutathione Sepharose 4B (Cytiva) resins, as described by the supplier. GST-tagged fusion proteins were eluted with 50 mM Tris–HCl, pH 8.5, 150 mM NaCl, 0.5% Triton X-100, 1 mM DTT, and 20 mM L-glutathione and then dialyzed against 10 mM Tris–HCl, pH 7.4, and 150 mM NaCl using a 7000 MWCO Slide-A-Lyzer Dialysis Cassette (Thermo Fisher Scientific). His-tagged proteins were eluted with the buffer containing 50 mM Tris–HCl, pH 7.6, 150 mM NaCl, 0.5% Triton X-100, 10% glycerol, 250 mM imidazole, and 1 mM MgCl_2_. The purity and concentration of the proteins were assessed by Coomassie brilliant blue staining of an SDS-PAGE gel of the eluent and the Bradford assay (Bio-Rad Laboratories, Hercules, CA, USA).

### In vitro kinase assay

Purified His-Pak4cat (55 nM) or His-Pak1cat (55 nM) was mixed with the purified protein of GST, GST-Inka1-iBox, or GST-Inka2-iBox (0.75 ng–400 ng), supplemented with 50 μM DTT and 20 μM ATP. The mixture was then incubated with 1 μg of AKT substrate II peptide (CKRPRAASFAE) as the Pak4 substrate (Promega) or 1 μg of GST-Raf13 as the Pak1 substrate with a total volume of 5 μL in a 384-well plate (PerkinElmer, Waltham, MA, USA) at 30°C for 1 h. For the termination of kinase reaction and depletion of the remaining ATP, 5 μL ADP-Glo reagent (Promega) was added to each reaction mixture and incubated for 40 min at 25°C. Then, a 10 μL kinase detection reagent (Promega) was added and incubated for 1 h at 25°C to convert the generated ADP to ATP. The amount of produced ATP was measured as a luminescence signal of luciferase activity using the Nivo S luminometer (PerkinElmer). The luminescent signal is proportional to the ADP concentration generated and reflects the Pak activity.

### Statistical analyses

Statistical analyses were performed using the R package version 3.4.2 and Prism9 version 9.1.2. All numerical data are expressed as mean ± SD. One-way ANOVA followed by Holm-Sidak’s multiple comparisons test or Weltch’s t-test with Holm– Bonferroni correction was used in multiple-group comparisons. Holm-Sidak’s multiple comparisons test was used to assess the significance of the actin polymerization ratio, luminescence, luciferase activity, number of dendritic spines, pCofilin, and pLIMK level between different groups, and qRT-PCR of neurons. Welch’s *t*-test was used to assess the significance of the LV area, CC thickness, the number of PSD95^+^ synapses, and qRT-PCR of MEFs. The Paired t-test was used to compare the pCofilin and pLIMK levels between different SN groups. The chi-squared test was used to compare the number of TUNEL^+^ cells and the cell morphology. *, P < 0.05; **, P < 0.01; ***, P < 0.001 were considered statistically significant. LV area and CC thickness were measured using ImageJ version 1.53a and normalized by the values of the control littermate. The number of apical and basal dendritic spines of the pyramidal neuron in cortical layer V were manually counted using cellSens imaging software (Olympus).

## Acknowledgment

The Inka2 (fam212b) mouse strain used for this research project was created from ES cell clones obtained from KOMP Repository (www.komp.org). The authors would like to thank Dr. Yohendran Baskaran and Dr. Ed Manser (Institute of Molecular and Cell Biology (A*STAR), Singapore) for sharing the Flag-Pak4cat, His-Pak4cat, GST-Inka1-iBox, GST-Inka2-iBox, GST-Raf13 plasmids. We would like to thank Mr. T. Yumoto and Mr. K. Otsu for their technical assistance. We also would like to thank Editage (www.editage.com) for English language editing. This work was funded by the Japan Society for the Promotion of Science grants-in-aid (KAKENHI) grant numbers 26430042 (to S.S.) and 19K06931 (to S.S.) and by Waseda University Grants for Special Research Projects 2017K-301 (to S.S.) and 2020C-372 (to S.S.).

## Author Contributions

Study concept and design: S.Y. and S.S. Acquisition of data: S.Y., T.M., A.T., and S.S. Analysis and interpretation of data: S.Y. and S.S. Drafting of the manuscript: S.Y. and S.S. Obtained funding: S.S.

## References

1. Ahmed, T., Shea, K., Masters, J.R., Jones, G.E., and Wells, C.M. (2008). A PAK4– LIMK1 pathway drives prostate cancer cell migration downstream of HGF. Cellular Signalling 20, 1320–1328.

2. Akiyama, H., Iwasaki, Y., Yamada, S., Kamiguchi, H., and Sakakibara, S. (2020). Control of cell migration by the novel protein phosphatase-2A interacting protein inka2. Cell and Tissue Research 380, 527–537.

3. Baskaran, Y., Ang, K.C., Anekal, P.V., Chan, W.L., Grimes, J.M., Manser, E., and Robinson, R.C. (2015). An in cellulo-derived structure of PAK4 in complex with its inhibitor Inka1. Nature Communications 6, 1–11.

4. Baskaran, Y., Ng, Y.W., Selamat, W., Ling, F.T.P., and Manser, E. (2012). Group I and II mammalian PAKs have different modes of activation by Cdc42. EMBO Reports 13, 653–659.

5. Baskaran, Y., Tay, F.P.-L., Ng, E.Y.W., Swa, C.L.F., Wee, S., Gunaratne, J., and Manser, E. (2021). Proximity proteomics identifies PAK4 as a component of Afadin– Nectin junctions. Nature Communications 12, 1–18.

6. Basu, S., and Lamprecht, R. (2018). The role of actin cytoskeleton in dendritic spines in the maintenance of long-term memory. Frontiers in Molecular Neuroscience 11, 143.

7. Borovac, J., Bosch, M., and Okamoto, K. (2018). Regulation of actin dynamics during structural plasticity of dendritic spines: Signaling messengers and actin-binding proteins. Molecular and Cellular Neuroscience 91, 122–130.

8. Chen, G.-L., Li, R., Chen, X.-X., Wang, J., Cao, S., Song, R., Zhao, M.-C., Li, L.-M., Hannemmann, N., and Schett, G. (2021). Fra-2/AP-1 regulates melanoma cell metastasis by downregulating Fam212b. Cell Death & Differentiation 28, 1364–1378.

9. Chen, L.Y., Rex, C.S., Babayan, A.H., Kramár, E.A., Lynch, G., Gall, C.M., and Lauterborn, J.C. (2010). Physiological activation of synaptic Rac> PAK (p-21 activated kinase) signaling is defective in a mouse model of fragile X syndrome. Journal of Neuroscience 30, 10977–10984.

10. Civiero, L., and Greggio, E. (2018). PAKs in the brain: Function and dysfunction. Biochimica et Biophysica Acta (BBA)-Molecular Basis of Disease 1864, 444–453.

11. Dan, C., Kelly, A., Bernard, O., and Minden, A. (2001). Cytoskeletal changes regulated by the PAK4 serine/threonine kinase are mediated by LIM kinase 1 and cofilin. Journal of Biological Chemistry 276, 32115–32121.

12. Dan, C., Nath, N., Liberto, M., and Minden, A. (2002). PAK5, a new brain-specific kinase, promotes neurite outgrowth in N1E-115 cells. Molecular and Cellular Biology 22, 567–577.

13. Darnell, J.C., Jensen, K.B., Jin, P., Brown, V., Warren, S.T., and Darnell, R.B. (2001). Fragile X mental retardation protein targets G quartet mRNAs important for neuronal function. Cell 107, 489–499.

14. Davis, J.K., and Broadie, K. (2017). Multifarious functions of the fragile X mental retardation protein. Trends in Genetics 33, 703–714.

15. del Mar Maldonado, M., and Dharmawardhane, S. (2018). Targeting rac and Cdc42 GTPases in cancer. Cancer Research 78, 3101–3111.

16. Dolan, B.M., Duron, S.G., Campbell, D.A., Vollrath, B., Rao, B.S., Ko, H.-Y., Lin, G.G., Govindarajan, A., Choi, S.-Y., and Tonegawa, S. (2013). Rescue of fragile X syndrome phenotypes in Fmr1 KO mice by the small-molecule PAK inhibitor FRAX486. Proceedings of the National Academy of Sciences 110, 5671–5676.

17. Frank, A.C., Huang, S., Zhou, M., Gdalyahu, A., Kastellakis, G., Silva, T.K., Lu, E., Wen, X., Poirazi, P., and Trachtenberg, J.T. (2018). Hotspots of dendritic spine turnover facilitate clustered spine addition and learning and memory. Nature Communications 9, 1–11.

18. Ghosh, M., Song, X., Mouneimne, G., Sidani, M., Lawrence, D.S., and Condeelis, J.S. (2004). Cofilin promotes actin polymerization and defines the direction of cell motility. Science 304, 743–746.

19. Hansen, A.H., Duellberg, C., Mieck, C., Loose, M., and Hippenmeyer, S. (2017). Cell polarity in cerebral cortex development—cellular architecture shaped by biochemical networks. Frontiers in Cellular Neuroscience 11, 176.

20. Hayashi, K., Ohshima, T., Hashimoto, M., and Mikoshiba, K. (2007). Pak1 regulates dendritic branching and spine formation. Developmental Neurobiology 67, 655–669.

21. Hayashi-Takagi, A., Araki, Y., Nakamura, M., Vollrath, B., Duron, S.G., Yan, Z., Kasai, H., Huganir, R.L., Campbell, D.A., and Sawa, A. (2014). PAKs inhibitors ameliorate schizophrenia-associated dendritic spine deterioration in vitro and in vivo during late adolescence. Proceedings of the National Academy of Sciences 111, 6461–6466.

22. Hayashi-Takagi, A., Takaki, M., Graziane, N., Seshadri, S., Murdoch, H., Dunlop, A.J., Makino, Y., Seshadri, A.J., Ishizuka, K., and Srivastava, D.P. (2010). Disrupted-in-Schizophrenia 1 (DISC1) regulates spines of the glutamate synapse via Rac1. Nature Neuroscience 13, 327–332.

23. Iwasaki, Y., Yumoto, T., and Sakakibara, S. (2015). Expression profiles of inka2 in the murine nervous system. Gene Expression Patterns 19, 83–97.

24. Liu, Y.Y., Tanikawa, C., Ueda, K., and Matsuda, K. (2019). INKA2, a novel p53 target that interacts with the serine/threonine kinase PAK4. International Journal of Oncology 54, 1907–1920.

25. Lupas, A., Van Dyke, M., and Stock, J. (1991). Predicting coiled coils from protein sequences. Science 252, 1162–1164.

26. Matsumura, H., Hasuwa, H., Inoue, N., Ikawa, M., and Okabe, M. (2004). Lineage-specific cell disruption in living mice by Cre-mediated expression of diphtheria toxin A chain. Biochemical and Biophysical Research Communications 321, 275–279.

27. McDonnell, A.V., Jiang, T., Keating, A.E., and Berger, B. (2006). Paircoil2: improved prediction of coiled coils from sequence. Bioinformatics 22, 356–358.

28. Meng, Y., Zhang, Y., Tregoubov, V., Janus, C., Cruz, L., Jackson, M., Lu, W.-Y., MacDonald, J.F., Wang, J.Y., and Falls, D.L. (2002). Abnormal spine morphology and enhanced LTP in LIMK-1 knockout mice. Neuron 35, 121–133.

29. Murray, B.W., Guo, C., Piraino, J., Westwick, J.K., Zhang, C., Lamerdin, J., Dagostino, E., Knighton, D., Loi, C.-M., and Zager, M. (2010). Small-molecule p21-activated kinase inhibitor PF-3758309 is a potent inhibitor of oncogenic signaling and tumor growth. Proceedings of the National Academy of Sciences 107, 9446–9451.

30. Nekrasova, T., Jobes, M.L., Ting, J.H., Wagner, G.C., and Minden, A. (2008). Targeted disruption of the Pak5 and Pak6 genes in mice leads to deficits in learning and locomotion. Developmental Biology 322, 95–108.

31. Ng, Y.-W., Raghunathan, D., Chan, P.M., Baskaran, Y., Smith, D.J., Lee, C.-H., Verma, C., and Manser, E. (2010). Why an A-loop phospho-mimetic fails to activate PAK1: understanding an inaccessible kinase state by molecular dynamics simulations. Structure 18, 879–890.

32. Nishiyama, J. (2019). Plasticity of dendritic spines: Molecular function and dysfunction in neurodevelopmental disorders. Psychiatry and Clinical Neurosciences 73, 541–550.

33. Pyronneau, A., He, Q., Hwang, J.-Y., Porch, M., Contractor, A., and Zukin, R.S. (2017). Aberrant Rac1-cofilin signaling mediates defects in dendritic spines, synaptic function, and sensory perception in fragile X syndrome. Science Signaling 10, eaan0852.

34. Qu, J., Cammarano, M.S., Shi, Q., Ha, K.C., de Lanerolle, P., and Minden, A. (2001). Activated PAK4 regulates cell adhesion and anchorage-independent growth. Molecular and Cellular Biology 21, 3523–3533.

35. Qu, J., Li, X., Novitch, B.G., Zheng, Y., Kohn, M., Xie, J.-M., Kozinn, S., Bronson, R., Beg, A.A., and Minden, A. (2003). PAK4 kinase is essential for embryonic viability and for proper neuronal development. Molecular and Cellular Biology 23, 7122–7133.

36. Reid, B.S., Sargent, T.D., and Williams, T. (2010). Generation and characterization of a novel neural crest marker allele, Inka1-LacZ, reveals a role for Inka1 in mouse neural tube closure. Developmental Dynamics 239, 1188-1196.

37. Rose, A., and Meier, I. (2004). Scaffolds, levers, rods and springs: diverse cellular functions of long coiled-coil proteins. Cellular and Molecular Life Sciences 61, 1996–2009.

38. Sells, M.A., Knaus, U.G., Bagrodia, S., Ambrose, D.M., Bokoch, G.M., and Chernoff, J. (1997). Human p21-activated kinase (Pak1) regulates actin organization in mammalian cells. Current Biology 7, 202–210.

39. Skarnes, W.C., Rosen, B., West, A.P., Koutsourakis, M., Bushell, W., Iyer, V., Mujica, A.O., Thomas, M., Harrow, J., and Cox, T. (2011). A conditional knockout resource for the genome-wide study of mouse gene function. Nature 474, 337–342.

40. Stricker, J., Falzone, T., and Gardel, M.L. (2010). Mechanics of the F-actin cytoskeleton. Journal of Biomechanics 43, 9–14.

41. Tabusa, H., Brooks, T., and Massey, A.J. (2013). Knockdown of PAK4 or PAK1 inhibits the proliferation of mutant KRAS colon cancer cells independently of RAF/MEK/ERK and PI3K/AKT signaling. Molecular Cancer Research 11, 109–121.

42. Tanabe, H., Goldstein, J., Yang, M., and Inouye, M. (1992). Identification of the promoter region of the Escherichia coli major cold shock gene, cspA. Journal of Bacteriology 174, 3867–3873.

43. Tian, Y., Lei, L., and Minden, A. (2011). A key role for Pak4 in proliferation and differentiation of neural progenitor cells. Developmental Biology 353, 206–216.

44. Villasana, L.E., Klann, E., and Tejada-Simon, M.V. (2006). Rapid isolation of synaptoneurosomes and post-synaptic densities from adult mouse hippocampus. Journal of Neuroscience Methods 158, 30–36.

45. Wiens, K.M., Lin, H., and Liao, D. (2005). Rac1 induces the clustering of AMPA receptors during spinogenesis. Journal of Neuroscience 25, 10627–10636.

46. Yamada, S., and Sakakibara, S. (2018). Expression profile of the STAND protein Nwd1 in the developing and mature mouse central nervous system. Journal of Comparative Neurology 526, 2099–2114.

47. Yamada, S., Sato, A., Ishihara, N., Akiyama, H., and Sakakibara, S. (2021). Drp1 SUMO/deSUMOylation by Senp5 isoforms influences ER tubulation and mitochondrial dynamics to regulate brain development. iScience 24, 103484.

48. Yamada, S., Sato, A., and Sakakibara, S. (2020). Nwd1 regulates neuronal differentiation and migration through purinosome formation in the developing cerebral cortex. iScience 23, 101058.

49. Zhang, H., Webb, D.J., Asmussen, H., Niu, S., and Horwitz, A.F. (2005). A GIT1/PIX/Rac/PAK signaling module regulates spine morphogenesis and synapse formation through MLC. Journal of Neuroscience 25, 3379–3388.

50. Zhang, K., Wang, Y., Fan, T., Zeng, C., and Sun, Z.S. (2022). The p21-activated kinases in neural cytoskeletal remodeling and related neurological disorders. Protein & Cell 13, 6–25.

51. Zhao, M., Spiess, M., Johansson, H.J., Olofsson, H., Hu, J., Lehtiö, J., and Strömblad, S. (2017). Identification of the PAK4 interactome reveals PAK4 phosphorylation of N-WASP and promotion of Arp2/3-dependent actin polymerization. Oncotarget 8, 77061.

