## Supplemental information for "Inka2, a novel Pak4 inhibitor, regulates actin dynamics in neuronal development"

**Table S1. The primers for genotyping of *Inka2* KO mice, qRT-PCR, and RIP analysis**

**Figure S1. Inka2 did not affect cell morphology in HeLa cells**

HeLa cells were transfected with control EGFP, EGFP-Inka2, EGFP-Inka2ΔiBox, or EGFP-Inka2ΔCC along with Flag-control. Cells were immunostained with anti-Flag antibody (magenta), and F-actin was stained with phalloidin (red). Scale bars, 20 μm.

**Figure S2. Co-expression of EGFP-Inka1 and Pak4cat often causes *in cellulo* crystallization of the protein complex.**

HEK293T cells were transfected with control EGFP-Inka2 or EGFP-Inka1 along with Flag-Pak4cat. Cells were immunostained with an anti-Flag antibody (red). Insets show a magnified view of the needle-shaped protein crystals growing within the individual cell. Simultaneous expression of EGFP-Inka1 and Pak4cat often caused *in cellulo* crystallization of the protein complex containing Inka1. Scale bars, 100 μm.

**Figure S3. F- and G-actin contents in HEK293T cells overexpressing Inka2 or Inka1 and purification of recombinant Pak4cat and Pak1cat**

(A, B) HEK293T cells were transfected with EGFP, EGFP-Inka1, and EGFP-Inka2, together with Flag-Pak4. Fractions of F-actin and G-actin were separated by ultracentrifugation from the cell lysates and quantified using immunoblotting with an anti-β-actin antibody. (B) Quantified analysis of the actin polymerization (F-actin/G-actin) ratio of five independent experiments. ns, not significant, *, $P < 0.05$; One-way ANOVA. Holm–Sidak's multiple comparisons test.

(C) Purification of Pak4cat and Pak1cat. Bacterially expressed and affinity-purified recombinant proteins of His-Pak4cat and Pros2-His-Pak1cat (PH-Pak1cat). The quality of the purified proteins is confirmed by SDS-PAGE and CBB staining.

**Figure S4. *Inka2* and *Inka1* mRNA expressions in the hippocampus**

(A–D) *In situ* hybridization of *Inka2* (A, B) and *Inka1* (C, D) mRNAs in the dentate gyrus of wild-type (A, C) or *Inka2*$^{-/-}$ adult mice (B, D). Scale bars, 50 μm in (A–D).

**Figure S5. Unchanged cell density in the *Inka2*$^{-/-}$ cerebral cortex, the normal polarization of Inka2$^{-/-}$ cortical neurons, and the validation of *Inka1* shRNAs.**

(A) The unaltered number of neurons in the *Inka2*$^{-/-}$ cerebral cortex. The cell density in cortical layer V of 12-month-old wild-type or *Inka2*$^{-/-}$ mice was counted. ns, not significant; Welch's t-tests.

(B) PCNs dissociated from wild-type or *Inka2*$^{-/-}$ embryonic cortices were cultured and then double-immunostained with anti-MAP2 and anti-SMI312 antibodies at 3 div. Scale bars, 20 μm.

(C) Validation of *Inka1* shRNAs. The non-targeting control or *Inka1* shRNA (shRNA #1 or shRNA #2) was transfected into HEK293T cells expressing EGFP-Inka1. Subsequently, each cell lysate was subjected to immunoblotting with anti-GFP and anti-α-tubulin antibodies.

Table S1

**Genotyping primer**
F1, forward: 5’-CATCAAGGAGCAATGTCACGGAAAGGAC-3’
R1, reverse: 5’-TCTGTCCACTACTCTTTGCAATTGTGAC-3’ (402bp)
F2, forward: 5’-GCTACCATTACCAGTTGGTCTGGTGTCA-3’
R2, reverse: 5’-TCAGCTCAGCCTGACCTGTGACAGACA-3’ (430bp)

**3'RACE primer**
*Inka2* GSP: 5’-GGACATGGACTGCTATCTGCGTCGCC-3’
*Inka2* NGSP: 5’-GCTATCTGCGTCGCCTCAAACAGGAGC-3’

**qRT-PCR primer**
*Inka2*, forward: 5’-ATCTGCGTCGCCTCAAACAGGA-3’
*Inka2*, reverse: 5’-TTCAGTTCCTGAAGTGCGCCCA-3’ (100bp)
*Inka1*, forward: 5’-TGTACAGCACCAGCACCAGCAT-3’
*Inka1*, reverse: 5’-TTTGGTTGCAGTGGGCCAGACA-3’ (124bp)
*β-actin*, forward: 5'-GGCTGTATTCCCCTCCATCG-3'
*β-actin*, reverse: 5'-CCAGTTGGTAACAATGCCATGT-3' (154bp)

**RIP primer**
*Inka2*, forward: 5’-GGATCCATTGAACCCAAAGGAG-3’
*Inka2*, reverse: 5’-AGTCTGCATTTTAGGTTCAGTGAGC-3’ (369bp)
*fmr1*, forward: 5’-ATTCCATGATGTGAGATTCCCACC-3’
*fmr1*, reverse: 5’-AGCAGGTTTGTTGGGATTAACAGA-3’ (235bp)
*map1b*, forward: 5’-TATTCCCAACCACAGCAACAGTAA-3’
*map1b*, reverse: 5’-CTGCTTGCTAAGACCATGATGTTG-3’ (267bp)
*gapdh*, forward: 5’-TAGACAAAATGGTGAAGGTCGGT-3’
*gapdh*, reverse: 5’-GTTCACACCCATCACAAACATGG-3’ (410 bp)

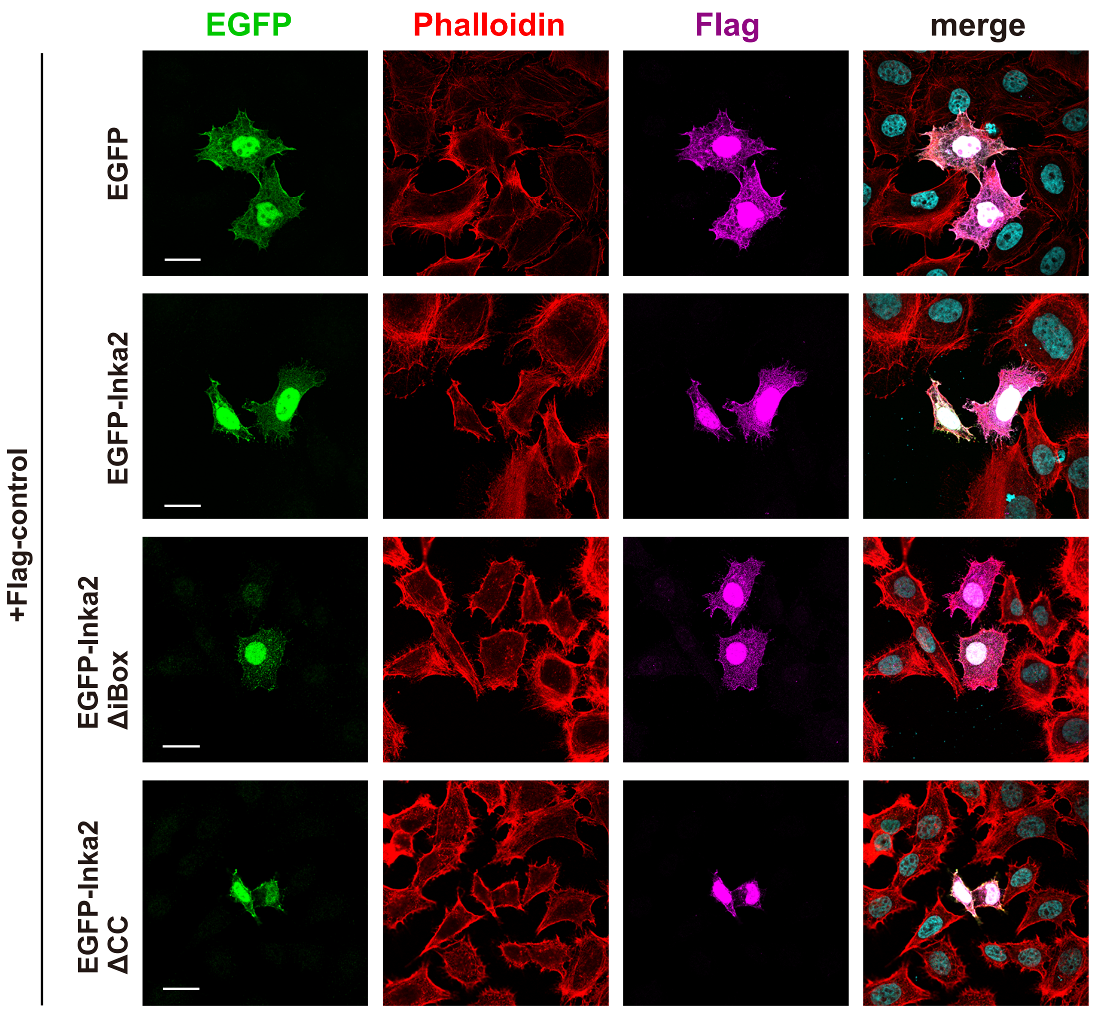
EGFP
Phalloidin
Flag
merge
+Flag-control
EGFP
EGFP-Inka2
EGFP-Inka2 ΔiBox
EGFP-Inka2 ΔCC

Figure S1

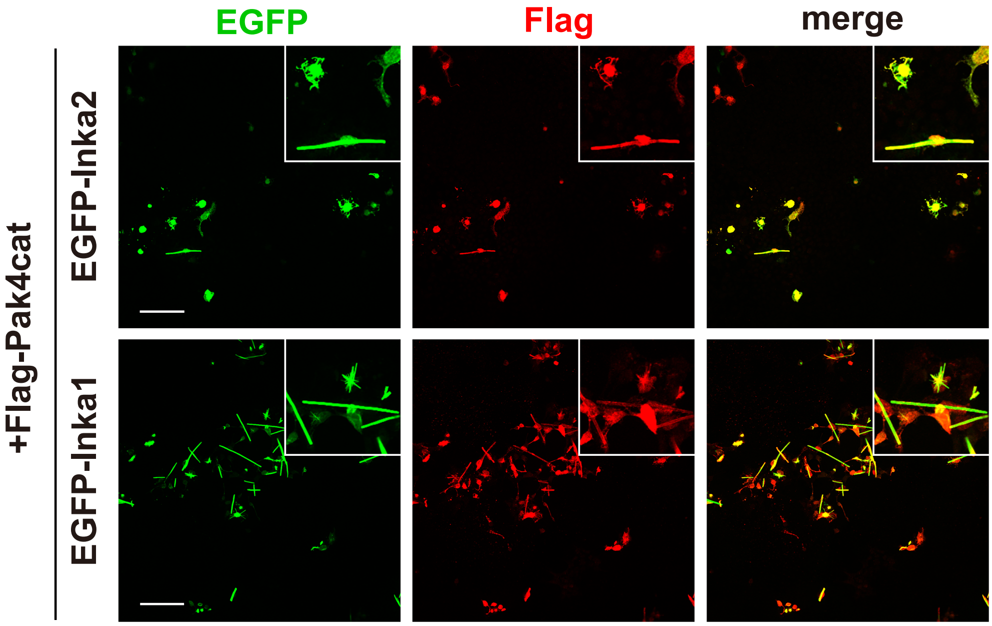


Figure S2

A

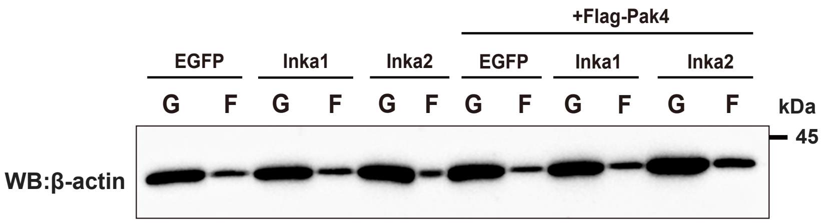


B

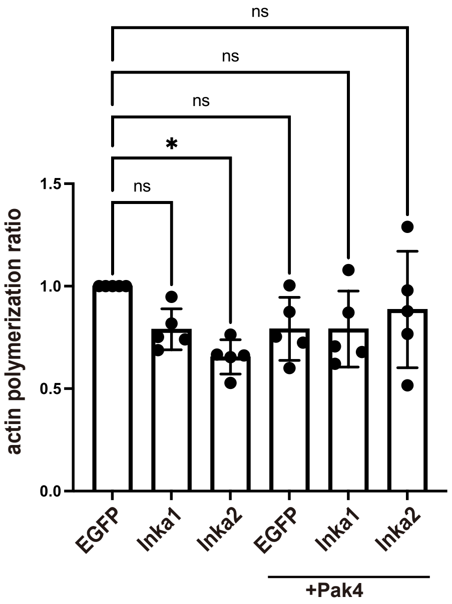


C

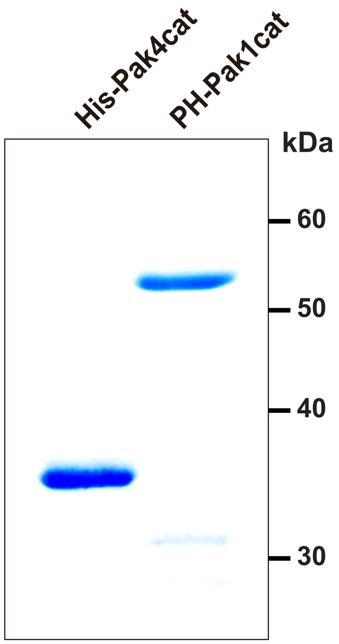


Figure S3

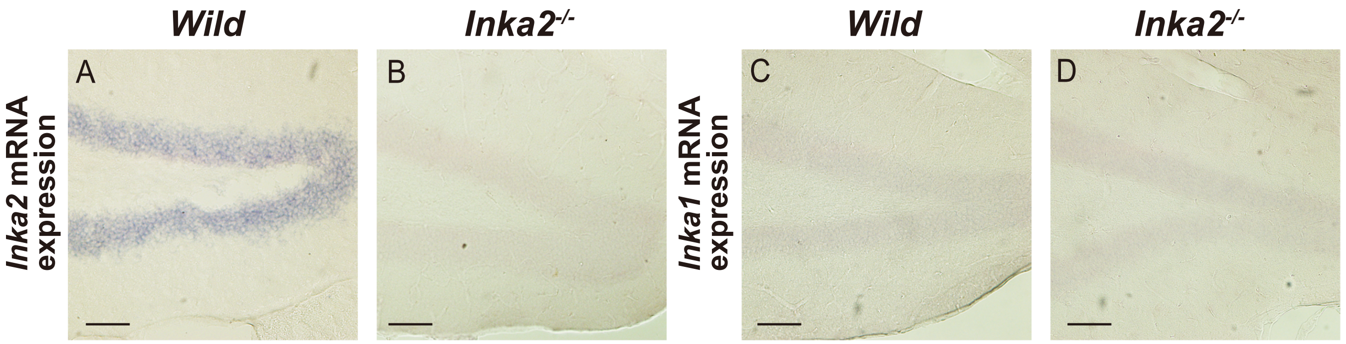
Wild
Inka2-/-
Wild
Inka2-/-
Inka2 mRNA expression
Inka1 mRNA expression
A
B
C
D

Figure S4

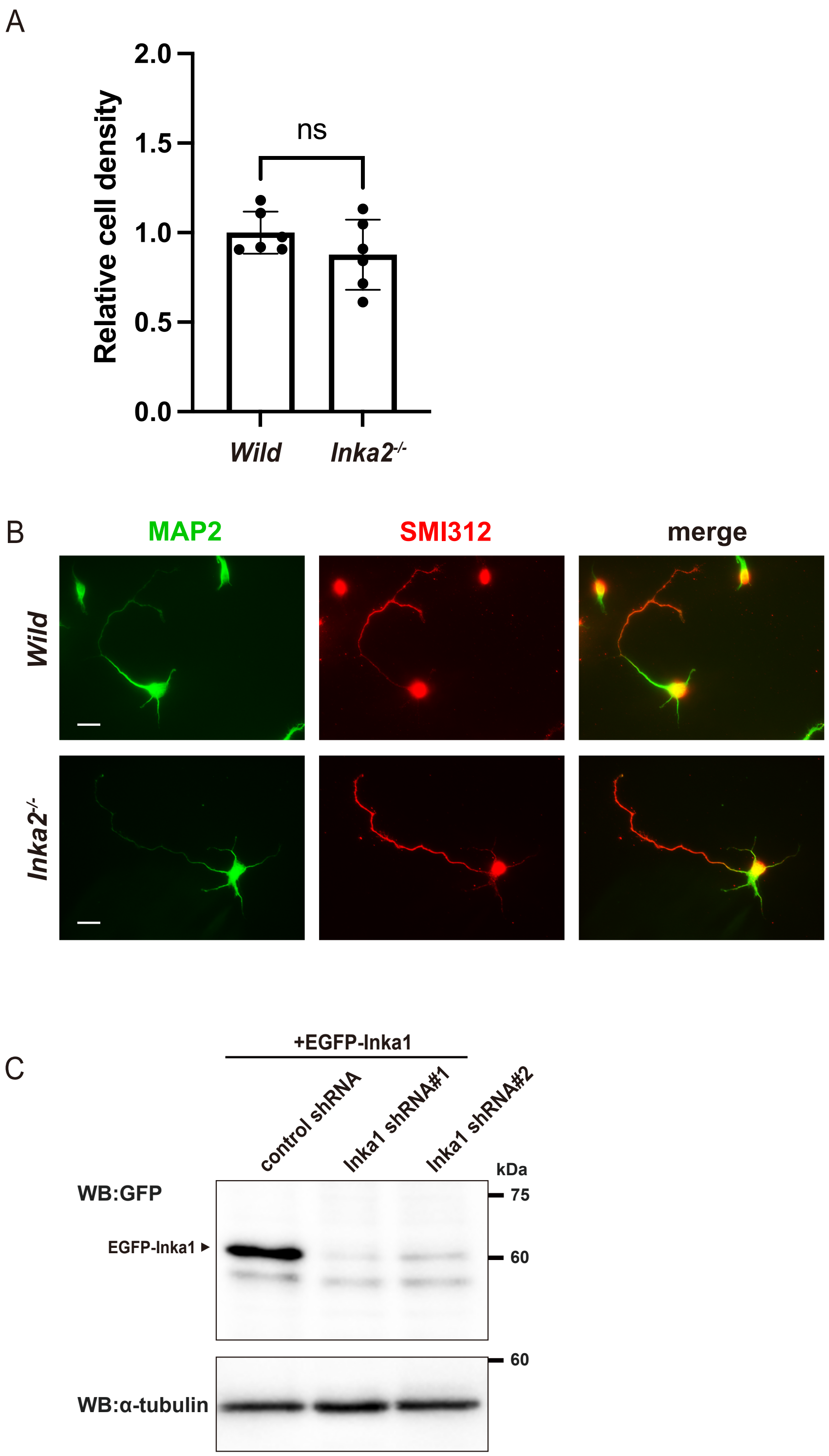
A
Relative cell density
2.0
1.5
1.0
0.5
0.0
ns
Wild
Inka2-/-
B
MAP2
SMI312
merge
Wild
Inka2-/-
C
+EGFP-Inka1
control shRNA
Inka1 shRNA#1
Inka1 shRNA#2
kDa
WB:GFP
75
EGFP-Inka1
60
60
WB:α-tubulin

Figure S5
